# SPO11 dimerization controls meiotic DNA double-strand break formation

**DOI:** 10.1101/2024.11.20.624454

**Authors:** Cédric Oger, Corentin Claeys Bouuaert

## Abstract

SPO11 initiates meiotic recombination through the induction of programmed DNA double-strand breaks (DSBs), but this catalytic activity had never been reconstituted *in vitro*. Here, using *Mus musculus* SPO11, we report a biochemical system that recapitulates all the hallmarks of meiotic DSB formation. We show that SPO11 catalyzes break formation in the absence of any partners and remains covalently attached to the 5′ broken strands. We find that target site selection by SPO11 is influenced by the sequence, bendability and topology of the DNA substrate, and provide evidence that SPO11 can reseal single-strand DNA breaks. In addition, we show that SPO11 is monomeric in solution and that cleavage requires dimerization to reconstitute two hybrid active sites. SPO11 and its partner TOP6BL form a 1:1 complex that catalyzes DNA cleavage with a similar activity to SPO11 alone. However, the complex binds DNA ends with higher affinity, suggesting a potential role post-cleavage. We propose a model where additional partners of SPO11 required for DSB formation *in vivo* assemble biomolecular condensates that recruit SPO11-TOP6BL, enabling dimerization and cleavage. Our work establishes SPO11 dimerization as the fundamental mechanism that controls the induction of meiotic DSBs.

## Introduction

During meiosis, SPO11 induces the formation of DNA double-strand breaks (DSBs) that initiate recombination, which is required for the accurate segregation of homologous chromosomes and promotes genetic diversity^1-3^. Despite >25 years since the discovery of SPO11, its catalytic activity had never been reconstituted, hindering a detailed understanding of the mechanism and regulation of SPO11-mediated DSB formation.

SPO11 evolved from the DNA-cleavage (A) subunit of a heterotetrameric (A2B2) type IIB topoisomerase, Topo VI^2,4^. Like Topo VI, SPO11 cleaves DNA via a pair of nucleophilic attacks from an active site tyrosine to the DNA backbone, producing breaks with 2-nucleotide 5′-overhangs with the transesterase covalently attached to the 5′ DNA ends^5-7^. Each DNA strand is cleaved by a hybrid active site composed of the catalytic tyrosine from one subunit and a metal-ion binding pocket contributed by the second subunit^8,9^. While Topo VI modulates DNA topology via cycles of gate opening (DNA cleavage), strand passage, and gate closing, orchestrated by the ATP-dependent Topo VIB subunit^10^, SPO11 in contrast cleaves DNA without restoring the broken strands. Nevertheless, *in vivo*, SPO11’s cleavage activity depends on a Topo VIB-like subunit (TOP6BL) and a series of additional accessory factors^11^. In *Mus musculus*, these include meiosis-specific partners REC114, MEI4, MEI1, and IHO1 (RMMI)^12-15^. However, in the absence of a reconstituted system, the mechanism that controls SPO11 activity and the function of the partners has remained unknown.

Here, we show that mouse SPO11 has intrinsic DNA-cleavage activity *in vitro* and explore the factors that impact target site selection and cleavage. We find that SPO11 cleavage is inherently limited by its monomeric state and propose that SPO11 dimerization constitutes the fundamental mechanism that controls the initiation of meiotic recombination.

## Results

### Mouse SPO11 has intrinsic DNA-cleavage activity

To set up an *in vitro* system to study meiotic DSB formation, we purified *Mus musculus* SPO11 from baculovirus-infected insect cells (**Fig. 1a, b**). To our surprise, we observed a robust plasmid DNA-cleavage activity that coincided with the peak of MBP-tagged SPO11 protein eluting from an ion exchange column (**Fig. 1c, d**). This activity was abolished when the tandem active site tyrosines (Y137, Y138) found in SPO11 were mutated to phenylalanine (YYFF) (**Fig. 1e**). Cleavage depended on the presence of a divalent metal ion, with Mn^2+^ being more effective than Mg^2+^, and Ca^2+^ also supporting low levels of cleavage (**Fig. 1f**). Hence, mouse SPO11 has intrinsic double-strand DNA cleavage activity, even in the absence of any of the partners known to be required *in vivo*.

**Fig. 1:**
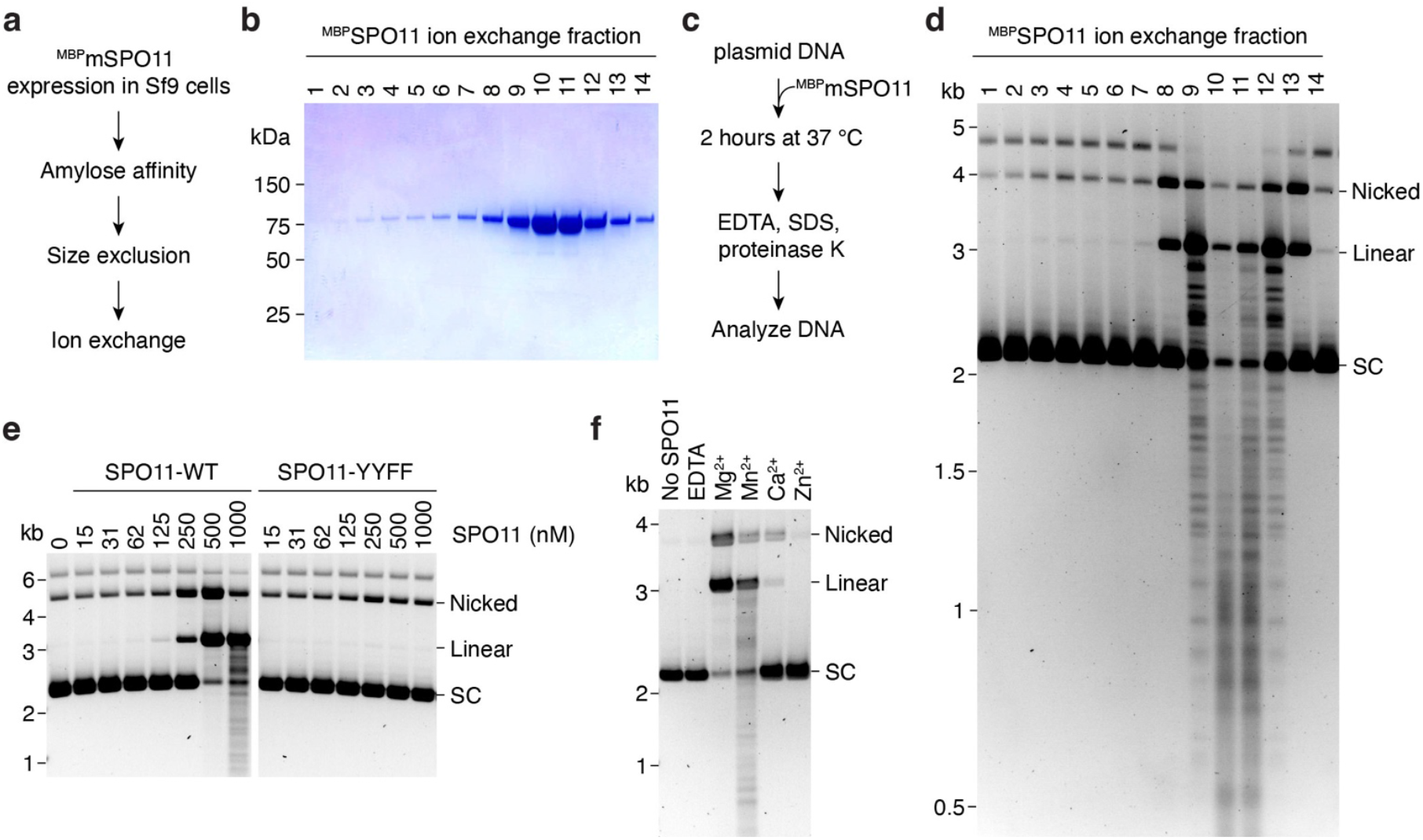
*In vitro* reconstitution of SPO11-dependent DNA double-strand break formation. **(a)** Purification scheme of mouse SPO11 protein. **(b)** SDS-PAGE analysis of ion exchange fractions of purified ^MBP^SPO11. **(c)** Scheme of the *in vitro* DNA cleavage assay. **(d)** Plasmid DNA cleavage analysis using fractions of SPO11 from panel b. SC, supercoiled plasmid. **(e)** Impact of the active site mutation YYFF (Y137F, Y138F) on the DNA-cleavage activity of SPO11. **(f)** Requirement for divalent metal ions on the DNA-cleavage activity of SPO11.

### SPO11 binds covalently to 5′ DNA strands

We verified that SPO11 remains covalently attached to the DNA breaks in four different ways.

First, we asked whether the linear cleavage product depended on deproteination of the sample prior to electrophoresis. Indeed, in the absence of proteinase K, the linear product was not detected and was instead converted to a smear due to the presence of covalently-bound denatured SPO11 proteins (**Fig. 2a**).

**Fig. 2:**
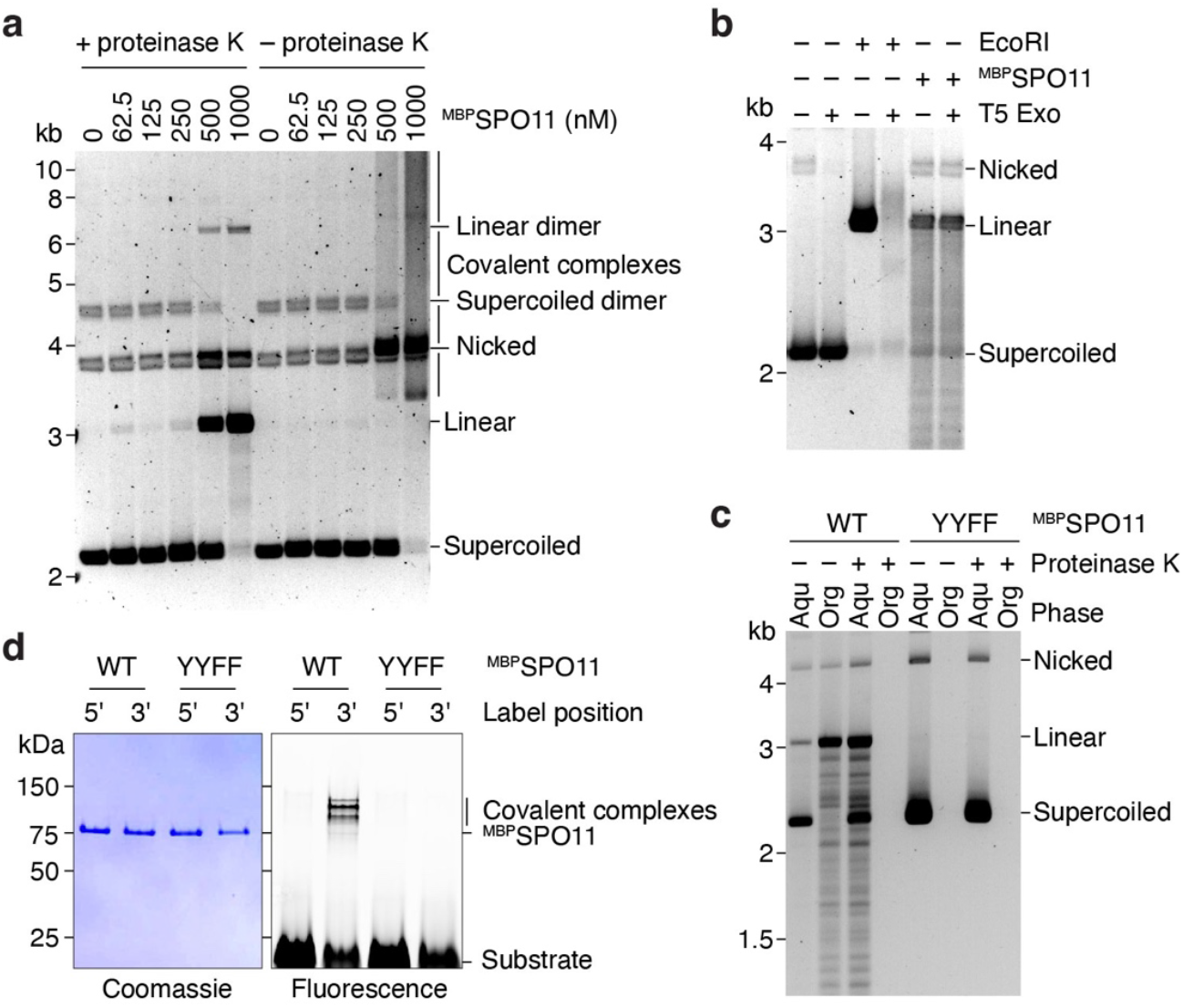
SPO11 is covalently bound to the 5′ DNA ends. **(a)** Agarose gel analysis of DNA cleavage products with or without proteinase K treatment prior to electrophoresis. All reactions were treated with SDS to eliminate non-covalent binding. **(b)** Analysis of the resistance of EcoRI or SPO11-dependent cleavage products to the 5′-3′ exonuclease T5 Exo. **(c)** Phenol-chloroform partitioning of DNA cleavage products with wild-type and mutant SPO11, with or without proteinase K treatment prior to phenol extraction. Aqu, aqueous phase; Org, organic phase. **(d)** SDS-PAGE analysis of covalent SPO11-DNA complexes from cleavage assays using wild-type or mutant SPO11 in the presence of 3′ or 5′ fluorescently labeled 80-bp substrates.

Second, we asked whether the broken DNA ends were protected from degradation by a 5′-3′ exonuclease. While a linear product generated by digestion with EcoRI was degraded in the presence of T5 exonuclease, the SPO11 cleavage products were resistant to exonuclease treatment (**Fig. 2b**).

Third, we tested whether phenol-chloroform partitioning leads to the enrichment of covalent protein-DNA adducts in the organic phase. As expected, cleaved DNA was detected in the organic phase in the presence of wild type SPO11, but not the YYFF mutant (**Fig. 2c**). If the samples were treated with proteinase K before phenol extraction, the cleavage products were absent from the organic phase and were instead found in the aqueous phase.

Fourth, we performed cleavage reactions with 5′ or 3′ fluorescently-labeled 80-bp duplex DNA substrates and separated covalent complexes by SDS-PAGE. Fluorescent signal of slightly lower electrophoretic mobility than ^MBP^SPO11 was detected with the 3′ labeled substrate, but not the 5′-labeled substrate, and was absent with the YYFF mutant (**Fig. 2d**). Hence, upon cleavage, SPO11 remains covalently bound to the 5′ broken DNA strands, as observed *in vivo*^5,6^.

### Cleavage pattern and substrate preferences

To establish whether SPO11 produces the expected staggered breaks with 2-nucleotide 5′-overhangs, we performed cleavage reactions with 5′-labeled 80-bp duplexes and separated cleavage products by denaturing gel electrophoresis. We detected sites of preferential cleavage, creating products of 16, 30, 48 and 63 nucleotides on the top strand and 15, 30, 48 and 62 nucleotides on the bottom strand (**Fig. 3a**). The position of the cleavage sites confirms the staggered cleavage pattern of SPO11 (**Fig. 3a**, bottom). Seven out of the eight preferred cleavage sites had a guanosine in position -3 with respect to the dyad axis, suggesting that base composition of the substrate influences SPO11 activity.

**Fig. 3:**
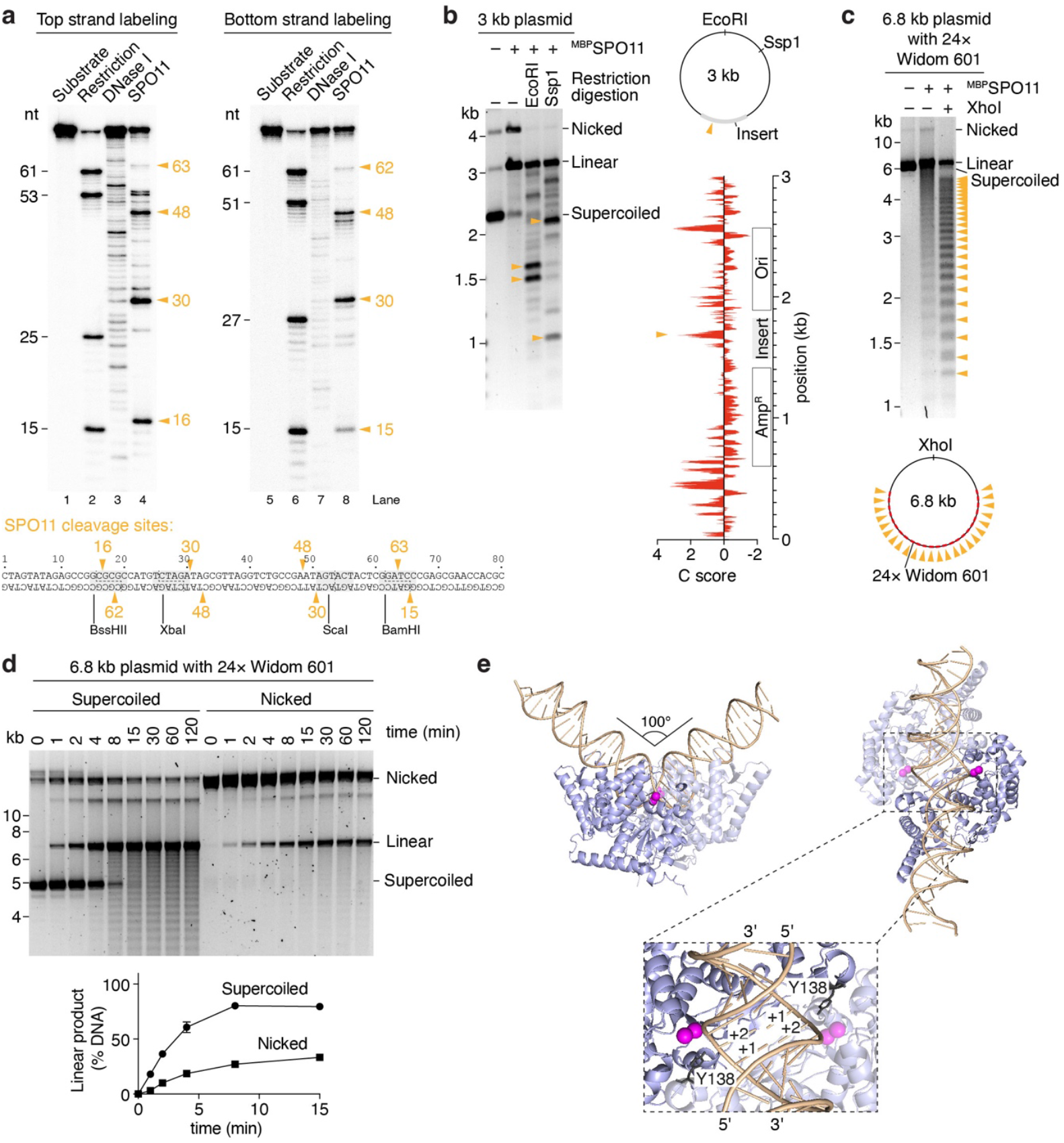
Cleavage pattern and substrate specificity. **(a)** Sequencing gel analysis of DNA cleavage reactions using 5′ radioactively labeled 80-bp substrates. Lanes 2 and 6 are produced by digestion of the substrate using restriction enzymes indicated below the gel. Lanes 3 and 7 are produced by partial digestion of the substrate with DNase I. SPO11 cleavage sites (lanes 4 and 8) are highlighted with orange arrowheads. **(b)** Agarose gel analysis of SPO11 cleavage sites on the standard plasmid substrate (pCCB959) using restriction digestion of SPO11 reaction products. Bottom right, cyclizability (C score) of the plasmid substrate, predicted by DNAcycP^20^. The position of the preferential cleavage site is indicated (arrowhead). **(c)** Analysis of SPO11 cleavage sites with a plasmid substrate that contains 24 copies of the Widom 601 sequence (pOC157). **(d)** Impact of DNA topology on the rate of SPO11-dependent cleavage. Quantifications show the mean and range from two independent experiments. **(e)** AlphaFold3 model of SPO11 dimer bound to a 40 bp duplex DNA substrate. Mg^2+^ ions are shown in magenta. The nucleotides that would form the 5′ overhang are labeled +1 and +2.

In mice, the distribution of SPO11-dependent breaks across the genome is largely determined by sequence-specific binding of the H3K4 methyltransferase PRDM9 that defines the position of ∼100-300 bp regions of elevated DSB activity (DSB hotspots)^16-19^. However, whether intrinsic substrate preferences of mouse SPO11 contribute to fine-scale selection of cleavage sites within hotspots is unknown.

In addition to potential contacts with DNA bases, substrate selection by SPO11 could be influenced by structural features of the double helix, e.g. DNA bending or unwinding. To explore the factors that impact SPO11 target site selection, we digested the products of our standard plasmid cleavage reaction with a restriction endonuclease. Agarose gel analysis revealed a non-random distribution of cleavage sites across the plasmid (**Fig. 3b**). A single prominent cleavage site mapped within a synthetic sequence cloned within the pUC-derived plasmid, while cleavage was much less efficient within the plasmid backbone. Using a DNA bendability prediction tool, DNAcycP^20^, we found that the preferential cleavage site corresponded to a peak of predicted DNA bendability (**Fig. 3b**, bottom right).

To test whether DNA bendability impacts SPO11 target site selection, we performed cleavage reactions using a plasmid that contains 24 repeats of the highly bendable Widom 601 sequence, a strong nucleosome binding sequence^21^. Restriction digestion of SPO11 cleavage reactions produced a strikingly periodic pattern, indicating that the Widom 601 sequences produce hotspots for DNA cleavage (**Fig. 3c**). To confirm this, we cloned 1, 3 and 6 copies of the Widom 601 sequence into a pUC19 plasmid and used this as PCR template to create linear substrates with a fluorophore at one extremity. Agarose gel analysis of SPO11 cleavage reactions allows us to map cleavage products along the linear substrate, which reveals a correlation between the cleavage sites, the position of Widom 601 sequences, and the predicted bendability of the substrate (**Extended Data Fig. 1**). Nevertheless, bendability alone as predicted by DNAcycP is not sufficient to account for the target site preference of SPO11. In addition, we cannot exclude that the Widom 601 sequence produces hotspots because of favorable base-specific contacts with SPO11, irrespective of bendability.

Next, we asked whether SPO11 cleavage is affected by the topology of the DNA substrate. Time-course analysis revealed that supercoiled plasmids are cleaved more efficiently than nicked plasmids (**Fig. 3d**). Hence, the cleavage activity of SPO11 is likely affected by a combination of inter-related factors including DNA sequence, bending, and topology.

Using AlphaFold3, we modeled the structure of a SPO11 dimer bound to a duplex DNA substrate (**Fig. 3e, Extended Data Fig. 2**). The model shows SPO11 poised for catalysis, with the active site tyrosine placed 3 Å from the correct phosphate groups to produce a break with 2-nucleotide 5′-overhangs. The substrate is bent with an angle of 100° with underwound DNA strands at the center of the complex, consistent with the observed preference of SPO11 for bendable and negatively supercoiled DNA.

### Double-strand DNA cleavage requires two hybrid active sites

All type II topoisomerases, including SPO11, are thought to cleave DNA using two composite active sites at the interface between two subunits^22-24^. The winged-helix domain of one subunit contributes the catalytic tyrosine, while the Toprim domain of the other subunit contributes metal-ion binding residues (**Fig. 4a**).

**Fig. 4:**
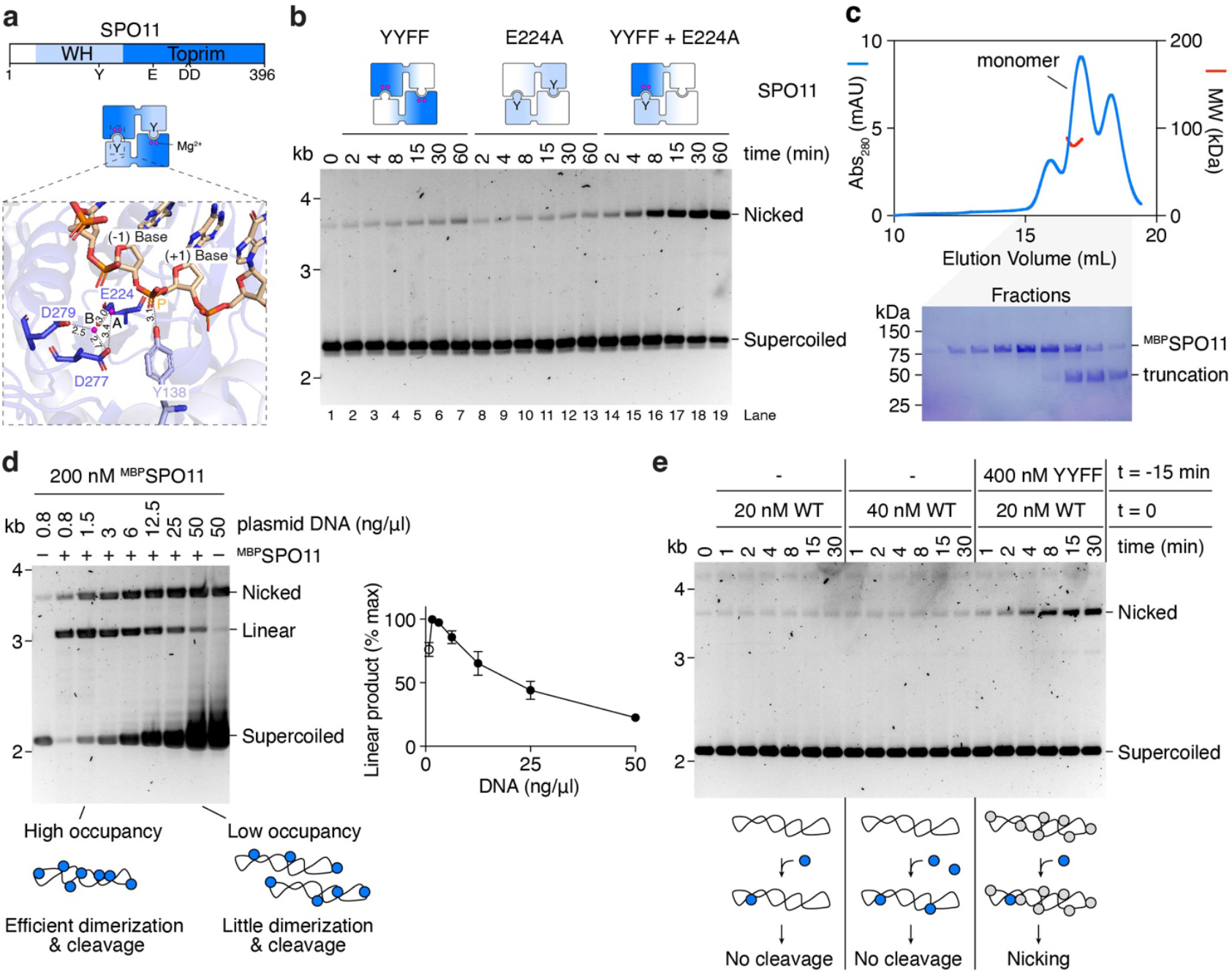
SPO11 cleavage requires dimerization. **(a)** Domain structure of SPO11 (top) and arrangement of the SPO11 dimer (center). Bottom, zoom of the composite active site within the AlphaFold3 model of a DNA-bound SPO11 dimer. Mg^2+^ ions are labeled A and B. Active site residues and their distances to metal ions and the scissile phosphate (p)are shown. WH, winged-helix domain. **(b)** Time-course analysis of SPO11 cleavage with mixtures of two catalytically-inactive mutants. **(c)** SEC-MALS analysis of ^MBP^SPO11 following amylose affinity purification. Blue traces are absorbance measurements at 280 nm from the size exclusion chromatography. Red traces are molar mass measurements across the peak. SDS-PAGE analyses of the corresponding fractions are shown. The molar mass of the leftmost peak could not be determined. The rightmost peak corresponds to a truncated fragment. **(d)** Titration of DNA at a constant concentration of SPO11. Reactions were stopped after 15 minutes. Quantifications show the mean and range from two independent experiments. At the lowest concentration (open circle) the substrate is limiting so the amount of linear product is not representative of total break levels. **(e)** Time-course analysis of SPO11 cleavage at the indicated concentrations of wild-type and catalytically-inactive SPO11. With 20 nM wild-type SPO11, the increased nicking activity observed in the presence of YYFF mutant cannot be explained by doubling the formation of cleavage-competent complexes, because no activity is detected at 40 nM wild type.

Most SPO11/Top6A homologs have two tandem tyrosines (Y137 and Y138 in mouse), except yeast that has a phenylalanine in the first position (**Extended Data Fig. 3a**). We confirmed by mutagenesis that Y138 is indeed the catalytic tyrosine in mouse SPO11 (**Extended Data Fig. 3b**).

The cleavage mechanism proposed for topoisomerases involves two metal ions, where metal ion A plays a direct role in catalysis, and metal ion B plays a structural role in stabilizing protein-DNA interactions^25,26^ (**Extended Data Fig. 3c**). The hybrid active site modeled by AlphaFold3 places Y138 in proximity to two metal ions, coordinated by residues E224, D277 and D279 (**Fig. 4a**). These residues are highly conserved in Top6A and SPO11 homologs (**Extended Data Fig. 3a**) and correspond to the active site residues previously identified in a crystal structure of *Methanocaldoccus jannaschii* Top6A^8^. We infer from sequence alignments that residues E224 and D277 coordinate metal ion A and residues D277 and D279 coordinate metal ion B. In yeast, E233 and D288 (equivalent to mouse E224 and D277) are essential for DSB formation, while D290 (equivalent to D279) is not^9^. Consistently, we found that mutating E224 to alanine abolishes the DNA-cleavage activity of mouse SPO11 (**Fig. 4b**, lanes 8-13).

To formally establish that cleavage involves hybrid active sites, we performed cleavage reactions using mixtures of two catalytically-inactive mutants, YYFF and E224A. The assembly of heterodimers should restore one functional active site per dimer, causing single-strand DNA breaks. As expected, we found that mixing the two inactive mutants leads to the formation of nicked products (**Fig. 4b**). Hence catalysis requires the concerted action of two SPO11 subunits to assemble hybrid active sites.

### Dimerization controls DNA cleavage

The mutant protein mixing experiment suggests either that purified SPO11 is monomeric in solution, like the yeast Spo11 core complex^27^ and *C. elegans* SPO-11^28^, or that dimers experience rapid subunit exchange. To determine the stoichiometry of SPO11, we subjected 4 µM ^MBP^SPO11 purified by amylose affinity to size exclusion chromatography followed by multi angle light scattering (SEC-MALS) (**Fig. 4c**). ^MBP^SPO11 produced a main peak that yielded an experimental molecular mass of 87.25 ± 1.5 kDa, consistent with a monomeric stoichiometry (expected MW: 87.4 kDa). This peak was preceded by a smaller one that may correspond to dimers, although the molecular weight could not be determined. Hence, the dissociation constant of SPO11 dimers is likely higher than 1-5 µM.

The predominantly monomeric stoichiometry of purified SPO11 accounts for its low intrinsic activity. Indeed, while SPO11 is active within a broad range of temperatures (optimum between 36°C and 42°C) and pH (6.5-8.5) (**Extended Data Fig. 4a, b**), cleavage is sensitive to the enzyme and substrate concentrations and requires a large excess of protein. For instance, with 12.5 nM (25 ng/µl) plasmid, no activity was detected under 60 nM SPO11, and 500 nM SPO11 was required to reach full conversion of supercoiled substrate into linear product within two hours (**Extended Data Fig. 4c**). When the substrate concentration was decreased, SPO11 cleaved DNA efficiently at concentrations as low as 50 nM, at least 10 to 100× lower than the expected KD (e.g. **Extended Data Fig. 4d**, lane 2). Hence, cleavage is not limited by the pool of soluble dimers but is instead a function of the protein-to-DNA ratio.

We reasoned that SPO11 binding to DNA would in effect increase its local concentration and perhaps allow dimerization directly on the DNA when the substrate occupancy is high enough. This would explain that increasing the DNA concentration inhibits cleavage (**Fig. 4d**), because reducing plasmid occupancy will reduce the chances of SPO11 dimerization.

If monomers meet on the substrate, DNA binding should be much more effective than cleavage. Indeed, gel shift analysis shows that SPO11 binds DNA at concentrations that do not support cleavage (**Extended Data Fig. 5**). Instead, at concentrations that do support cleavage, SPO11 binds so abundantly to the substrate that it provides effective protection against DNase I treatment (**Extended Data Fig. 6**). In addition, we found that cleavage is significantly more sensitive to salt than DNA binding (**Extended Data Fig. 7**), likely because a mild reduction of plasmid occupancy caused by increasing salt concentration strongly reduces the chances of dimerization. Finally, we found that preincubation of plasmid substrates with an excess of inactive SPO11 facilitates cleavage upon addition of low levels of wild-type protein, presumably because decreasing the search space on the substrate increases the rate of dimerization and cleavage (**Fig. 4e**).

Overall, these data indicate that SPO11 monomers bind efficiently to DNA and that, above a certain threshold, monomers meet on the substrate, allowing cleavage.

### SPO11 can reseal single-strand nicks

In conditions where SPO11 mostly cleaves a single DNA strand, for example in reactions that contain a mixture of wild type and YYFF mutant or YYFF and E224A mutants, we observe additional bands that migrate between the position of the supercoiled and nicked products (**Extended Data Fig. 8a, b**). Phenol-chloroform partitioning shows that these products are not covalently bound to SPO11, indicating that they are plasmid topoisomers (**Extended Data Fig. 8c**). Time-course analysis shows that these topoisomers accumulate more slowly than cleavage products (**Extended Data Fig. 8b**).

The formation of topoisomers implies that SPO11 can sometimes re-ligate broken DNA strands. The most likely scenario is that the topoisomers are caused by the separation of the two monomers when catalysis is stalled at the single-strand break, either accompanied by a dissociation of the inactive subunit from the DNA substrate or not (**Extended Data Fig. 8d**). This would result in the swiveling of the DNA around the intact phosphodiester bond. Re-assembly of the dimer interface would then provide an opportunity for reversal of the reaction and liberation of SPO11 from the substrate.

### The SPO11-TOP6BL complex

Mouse SPO11 forms a complex with the topoisomerase-derived TOP6BL subunit, which is required for DSB formation *in vivo*^11^. In Topo VI, the B subunit coordinates ATP-dependent dimerization of its GHKL domain with DNA cleavage by Top6A to control strand passage^10^. However, the role of TOP6BL in SPO11-dependent cleavage remains unclear.

To gain insight into the SPO11-TOP6BL complex, we used AlphaFold3 to model a 2:2 heterotetramer bound to a 40 bp duplex DNA (**Fig. 5b, Extended Data Fig. 2**). Similar to the SPO11-DNA complex presented above, the model shows SPO11-TOP6BL engaged with a bent DNA substrate, with the active site residues poised for cleavage. The complex showed high structural similarity to Topo VI, with TOP6BL presenting well-defined GHKL and transducer domains (**Extended Data Fig. 9**). However, Top6B has a helix two-turn helix (H2TH) motif located between the GHKL and transducer domains and some species have a structured C-terminal extension that are absent in TOP6BL. In addition, the ATP-binding site of Top6B is predicted to be degenerated in TOP6BL, consistent with previously reported models^11,29^.

**Fig. 5:**
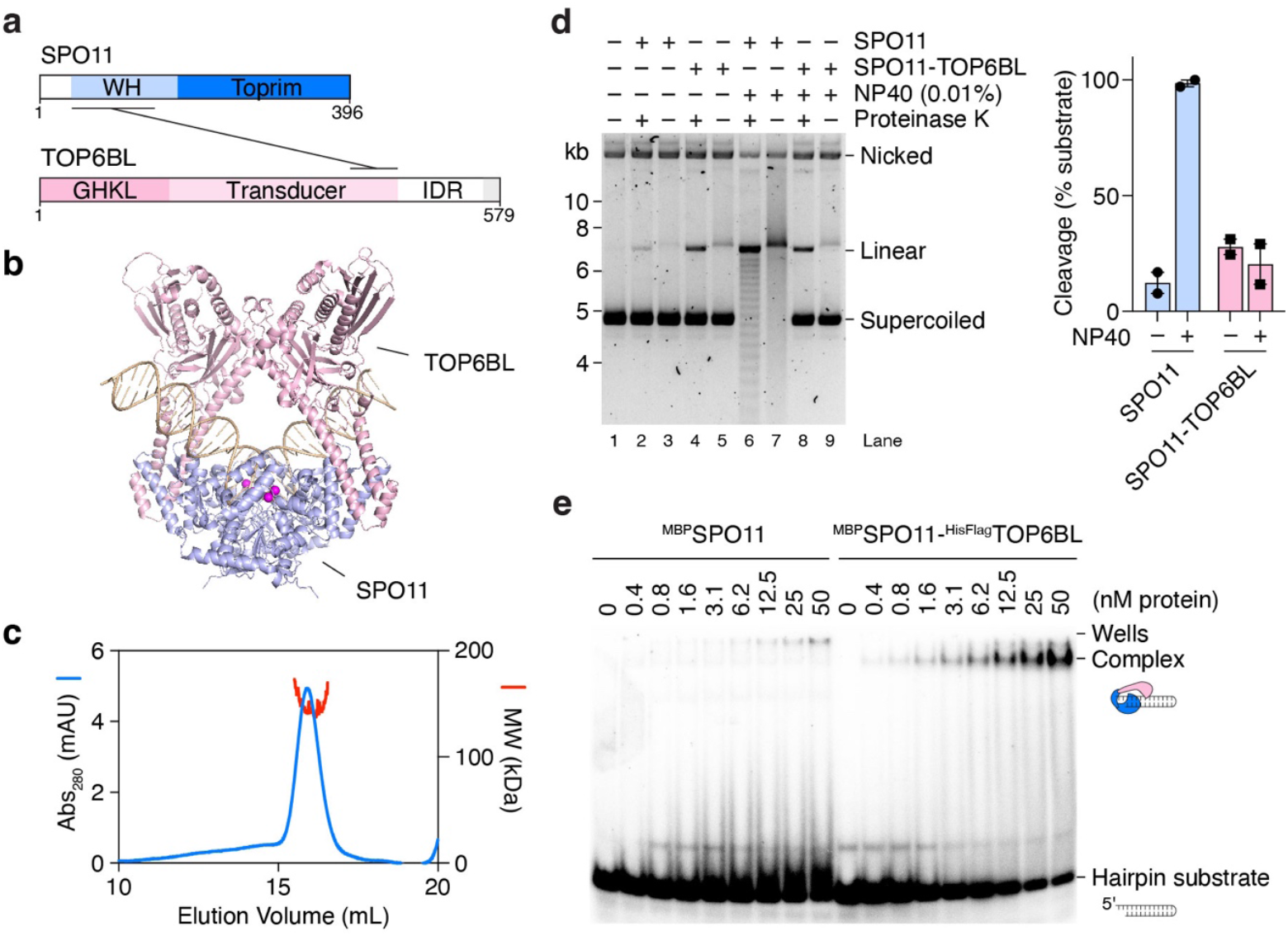
The SPO11-TOP6BL complex. (a)Domain structure of SPO11 and TOP6BL. IDR, intrinsically disordered region. The C-terminus of TOP6BL (grey) binds REC114^29^. **(b)** Alphafold3 model of the SPO11-TOP6BL heterotetramer bound to a 40 bp DNA substrate. The TOP6BL-IDR was omitted from the model. **(c)** SEC-MALS analysis of ^MBP^SPO11-^HisFlag^TOP6BL complexes. Blue traces are absorbance measurements at 280 nm from the size exclusion chromatography. Red traces are molar mass measurements across the peak. **(d)** Plasmid (pOC157) cleavage with 45 nM SPO11 or SPO11-TOP6BL complexes with or without 0.01% NP40. In lanes 3, 5, 7 and 9, proteinase K treatment was omitted prior to electrophoresis. Quantifications show individual data points, mean and range from two independent experiments. **(e)** Gel shift analysis of the binding of SPO11 and SPO11-TOP6BL complexes to a 25 bp hairpin substrate with a 2-nucleotide 5′-overhang.

To investigate the properties of SPO11-TOP6BL complexes, we purified ^MBP^SPO11-^HisFlag^TOP6BL from baculovirus-infected insect cells and analyzed complexes by SEC-MALS (**Fig. 5c**). This revealed an experimental molecular mass of 150.2 ± 2.2 kDa consistent with a 1:1 complex (expected MW: 157 kDa). Hence, just like SPO11, the cleavage activity of the SPO11-TOP6BL complex is likely to be limited by protein dimerization.

As anticipated, a plasmid cleavage assay revealed similar activities between SPO11 and SPO11-TOP6BL complexes, although this was context dependent. For instance, in our standard cleavage assay, at low protein concentrations, the SPO11-TOP6BL complex was slightly more active than SPO11 alone (**Fig. 5d**, lanes 2 and 4). However, in the presence of 0.01% NP40, the cleavage activity of SPO11 was strongly stimulated, while that of the SPO11-TOP6BL complex was not (lanes 6 and 8). We conclude that physicochemical parameters of the reaction impact differently the cleavage activity of SPO11 and SPO11-TOP6BL complexes, perhaps due to effects of TOP6BL on DNA binding. Indeed, purified mouse TOP6BL was recently shown to bind DNA^30^. In addition, the B subunit of Topo VI directly contacts DNA to coordinate substrate binding with ATP-dependent dimerization and catalysis^31^.

We previously showed that purified yeast Spo11 complexes, despite being catalytically inactive, bind with high affinity to DSBs through non-covalent interactions^32,33^. To investigate the end-binding activity of the mouse homologs, we performed gel shift analyses using a short hairpin substrate with a 2-nucleotide 5′-overhang at one extremity, which the yeast complex binds to with sub-nanomolar affinity. We found that the SPO11-TOP6BL complex binds efficiently to this substrate, while SPO11 alone does not (**Fig. 5e**). This confirms that TOP6BL affects the DNA-binding properties of SPO11 and supports a model where tight end-binding by SPO11-TOP6BL could impact DSB processing, as we previously proposed for the yeast complex^27,34^.

## Discussion

Here, we reconstituted the formation of meiotic DNA double-strand breaks *in vitro* using *Mus musculus* SPO11. We demonstrate that the *in vitro* assay recapitulates all of the features expected of a *bona fide* SPO11 cleavage assay: catalysis depends on the catalytic tyrosine Y138 and divalent metal ions; the protein remains covalently bound to the 5′-DNA strand; and each strand is cleaved by a composite active site assembled at the interface of two SPO11 monomers, with the two cleavages staggered to produce a 2-nucleotide 5′-overhang. In addition, we show that SPO11 cleavage site selection is driven by a mild sequence bias and may reflect in part a preference for bendable and underwound DNA, and found that SPO11 is able to reseal a broken strand when stuck in a single-strand nicked intermediate. SPO11 cleavage is inherently controlled by its monomeric state and occurs *in vitro* following dimerization on the DNA substrate. The accompanying paper by Zheng and colleagues reports similar findings focusing on the SPO11-TOP6BL complex. These results provide a framework to understand the mechanism and control of meiotic DNA double-strand break formation.

### A model for the formation of meiotic DNA double-strand breaks

In all eukaryotes where this has been investigated, SPO11 requires a cohort of accessory partners to catalyze break formation *in vivo*^3,34,35^. Our observation that SPO11 cleaves DNA *in vitro* in the absence of any additional factor constrains the putative functions of these partners, at least in mice. Instead, we show that an important intrinsic limit to SPO11 cleavage is dimerization.

In the *in vitro* assay, cleavage occurs at high protein/DNA ratios, which we interpret as allowing encounters of SPO11 monomers on the DNA substrate. The high DNA concentrations encountered *in vivo* will effectively titrate SPO11 monomers, preventing dimerization and cleavage. Hence, we suggest that a primary function of the partners is to control when and where SPO11 dimerizes.

In mice, the known essential partners of SPO11 are TOP6BL, REC114, MEI4, IHO1 and MEI1 (RMMI) (**Fig. 6a**)^11-15^. We have shown that SPO11-TOP6BL forms a 1:1 complex that has comparable cleavage activity to that of SPO11 alone, although this depended on the experimental conditions. In addition, we previously showed that the yeast orthologs of REC114, MEI4, IHO1 (Rec114, Mei4 and Mer2) assemble DNA-dependent condensates *in vitro* that recruit the yeast Spo11 core complex (Spo11, Ski8, Rec102, Rec104)^32^. Despite extreme sequence divergence, the structure of these proteins is widely conserved across eukaryotes, although their DNA-binding and condensation activities appear to vary quantitatively between organisms^36,37^.

**Fig. 6:**
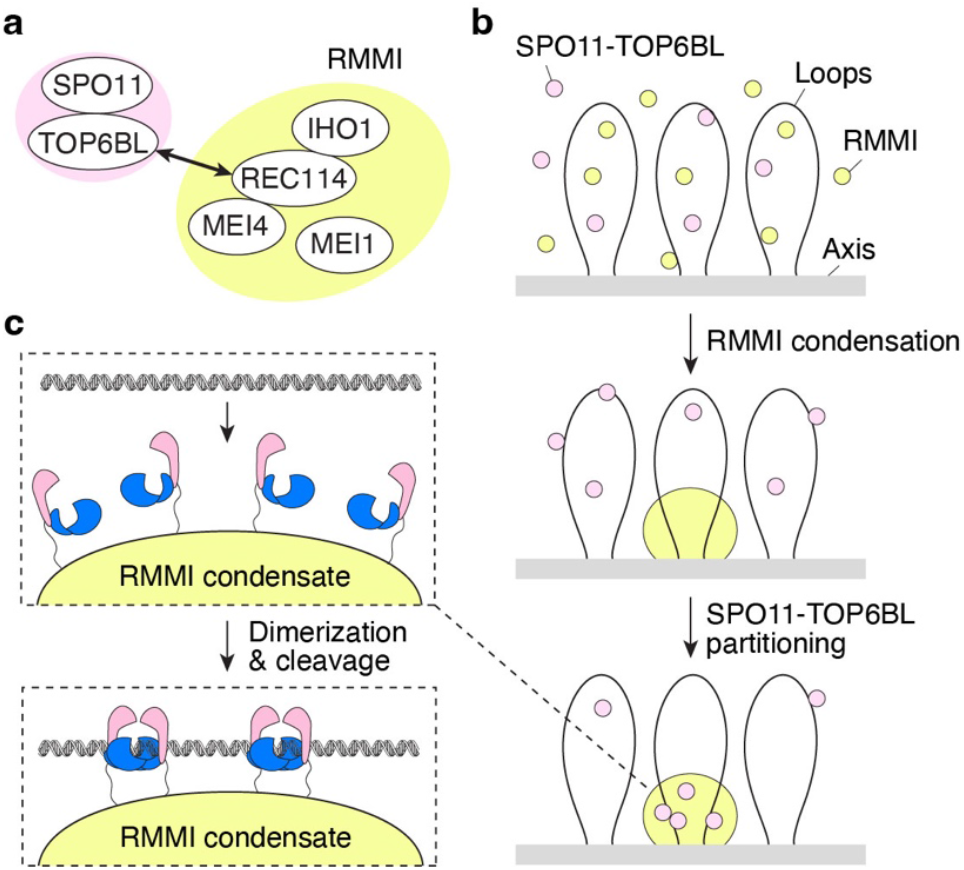
Model of meiotic DSB formation in mice. **(a)** Proteins essential for DSB formation in mice. **(b)** We propose that RMMI form condensates along the chromosome axis and recruit SPO11-TOP6BL complexes through an interaction between REC114 and the C-terminus of TOP6BL^29^. **(c)** The ensuing increase in local concentration of SPO11-TOP6BL complexes allows SPO11 dimerization and cleavage. Multiple dimers may assemble, leading to the formation of closely-spaced double DSBs^38,39^.

Based on these insights, we propose that the mouse RMMI proteins control SPO11 cleavage by assembling biomolecular condensates within which SPO11-TOP6BL is recruited (**Fig. 6b**). Recruitment of SPO11-TOP6BL complexes likely involves the interaction between a TOP6BL C-terminal α-helix that binds the pleckstrin homology (PH) fold of REC114^29^. Partitioning of SPO11-TOP6BL complexes within these condensates would allow them to reach a critical threshold required for dimerization and DSB formation.

In yeast and mice, SPO11 frequently catalyzes multiple closely-spaced DSBs, in particular in the absence of the DNA-damage response kinase Tel1/ATM^38-40^, which is consistent with local accumulation of SPO11 within condensates^32^. The distance between these coincident breaks ranges from about 30 nt to several kb, and the more closely-spaced events show a 10-bp periodicity, indicative of co-orientation of SPO11 dimers on the DNA substrate^38,39^. Our results suggest that, in addition to increasing the local concentration of SPO11, condensates function to co-orient SPO11 proteins to facilitate productive dimerization on the DNA substrate (**Fig. 6c**).

### Substrate site selection

In mice, the distribution of meiotic DSBs across the genome is primarily dictated by the H3K4 methyltransferase PRDM9 that binds specific sequence motifs through a fast-evolving zinc-finger array^16-18^. Here, we show that *in vitro* the cleavage of SPO11 is directly impacted by a combination of a mild sequence bias (preference for a G three nucleotides 5′ of the dyad axis) and probably a preference for bendable and/or underwound DNA. These factors are likely to influence the fine-scale selection of target sites within DSB hotspots and may lead to preferential induction of DSBs at sites that are under topological stress.

### Cleavage reversal and the dimer interface

We have shown that reaction conditions that accumulate single-strand cleavage products tend to generate covalently closed topoisomers, thereby revealing the ability of SPO11 to re-ligate a broken DNA strand. Partial relaxation of the plasmid before religation presumably requires the separation of the dimer interface, accompanied or not by the dissociation of the monomer that was not covalently bound to the broken strand. The swiveling of the DNA duplex around the intact strand that ensues is therefore a direct consequence of the weak dimer interface between SPO11 subunits. This is consistent with the monomeric state of the free protein and our model that dimerization controls DNA cleavage.

Given the weak dimer interface and the slow rate of the religation activity, we do not envision that SPO11 frequently reseals DSBs *in vivo*. Instead, DSB formation will most likely lead to rapid collapse of the dimer interface and separation of the two DNA ends. The ends may however remain in proximity through anchoring of SPO11-TOP6BL to RMMI condensates and loading of MRN complexes and their end-tethering activities^32,41^.

## Materials and Methods

### Preparation of expression vectors

Plasmids are listed in **Supplementary Table S1**. The sequences of the oligos are listed in **Supplementary Table S2**, and gBlocks (Integrated DNA Technologies) are listed in **Supplementary Table S3**.

Sequences coding for *M. musculus* SPO11 and TOP6BL were codon-optimized for expression in Sf9 cells and synthesized as gBlocks. The SPO11 gBlock was cloned into pFastBac1-MBP to yield pCCB630 (^MBP^SPO11). The TOP6BL gBlock was cloned into pFastBac1-HisFlag to yield pCCB628 (^HisFlag^mTOP6BL).

The SPO11-YYFF mutant was generated by QuikChange mutagenesis of pCCB630 using primers cb886 and cb887 to yield pCCB642 (^MBP^SPO11-YYFF). Other SPO11 active site mutants were generated by inverse PCR and self-ligation using template pCCB630. Primers and resulting plasmids are as follows: SPO11-Y137F (primers cb1577 and cb1578, plasmid pCCB1084), SPO11-Y138F (primers cb1579 and cb1580, plasmid pCCB1085), SPO11-E224A (primers cb1581 and cb1582, pCCB1086).

### Expression and purification of recombinant proteins

Viruses were produced using a Bac-to-Bac Baculovirus Expression System (Invitrogen) according to the manufacturer’s instructions. We infected 4×10^9^ *Spodoptera frugiperda* Sf9 cells (Gibco, Thermo Fisher) with viruses at a multiplicity of infection (MOI) of 2. Expression of ^MBP^SPO11 used viruses generated from pCCB630, ^MBP^SPO11-^HisFlag^TOP6BL used viruses generated from pCCB630 and pCCB628. After 72 hours infection, cells were collected, washed with phosphate buffer saline (PBS), frozen in dry ice and kept at -80°C until use. All purification steps were carried out at 0-4°C. Cell pellets were resuspended in 80 ml of lysis buffer (50 mM HEPES-NaOH pH 6.8, 1 mM DTT, 2 mM EDTA, protease inhibitor cocktail (Sigma-Aldrich, P8340) diluted 1:800, supplemented with 4 µM leupeptin, 5.8 µM pepstatin, 6.6 µM chymostatin and 1 mM phenylmethanesulfonyl fluoride (PMSF)) and then pooled in a beaker and mixed slowly with a stir bar for 20 minutes. 10% of ice-cold glycerol and 1 M NaCl were added to the cell lysate that was then centrifuged at 43,000 g for 25 min. The cleared extract was loaded onto 2 ml pre-equilibrated amylose resin (NEB). The column was washed extensively with amylose buffer (25 mM HEPES-NaOH pH 6.8, 1 M NaCl, 5% glycerol, 1 mM DTT, 2 mM EDTA) and eluted with buffer containing 10 mM maltose. Fractions containing protein were loaded on a HiLoad 16/600 Superdex 200 pg column preequilibrated with buffer containing 25 mM HEPES 6.8, 100 mM NaCl, 2 mM DTT, and 5 mM EDTA. The peak was collected and diluted 2-fold in buffer without salt, loaded on a Capto HiRes cation exchange column, and eluted with a 0.1-0.5 M NaCl gradient. Fractions containing purified proteins were pooled, aliquots were flash frozen in liquid nitrogen and stored at −80°C.

For the ^MBP^SPO11-^HisFlag^TOP6BL complex, Sf9 cells were lysed by bringing up salt concentration to 500 mM. The cleared extract was incubated 20 minutes with 2 ml pre-equilibrated Ni-NTA resin (Thermo Scientific) and washed extensively with a buffer containing 25 mM HEPES pH 7.5, 500 mM NaCl, 10% glycerol, 0.1 mM DTT and 40 mM imidazole. The complex was eluted with buffer containing 500 mM imidazole and loaded onto 2 ml of equilibrated amylose resin. The resin was washed with 25 mM HEPES 7.5, 500 mM NaCl, 10% glycerol, 1 mM DTT, 2 mM EDTA and the protein eluted with buffer containing 10 mM maltose. Fractions containing protein were loaded on a Superdex 200 column equilibrated in 25 mM HEPES pH 7.5, 400 mM NaCl, 10% glycerol, 2 mM DTT, 5 mM EDTA. The peak was collected, concentrated using 10 kDa Amicon centrifugal filters (Millipore), aliquoted, flash frozen in liquid nitrogen, and stored at −80°C.

### SEC-MALS

Light scattering data were collected using a Superdex 200 increase 10/300 GL Size Exclusion Chromatography (SEC) column connected to a AKTA Pure Chromatography System (Cytiva). The elution from SEC was monitored by a differential refractometer (Optilab, Wyatt), and a static and dynamic, multiangle laser light scattering (LS) detector (miniDAWN, Wyatt). The SEC–UV/LS/RI system was equilibrated in buffer 25 mM HEPES-NaOH pH 7.5, 500 mM NaCl, 10% glycerol, 5 mM EDTA, 2 mM DTT at a flow rate of 0.3 ml/min. The weight average molecular masses were determined across the entire elution profile at intervals of 0.5 seconds from static LS measurement using ASTRA software.

### Plasmid cleavage assays

Cleavage reactions (20 µl) were typically carried out with 250 nM ^MBP^SPO11, 5 ng/µl of a pUC19-derived 3-kb plasmid (pCCB959) in buffer containing 25 mM Tris pH 7.5, 5 % glycerol,

50 mM NaCl, 1 mM DTT, 0.1 mg/ml BSA, 5 mM MgCl_2_ and 1.5 mM MnCl_2_, unless stated otherwise. Reactions were incubated for 2 hours at 37°C, stopped with 50 mM EDTA and 1% SDS, and treated with 0.2 mg/ml proteinase K for 15-30 minutes at 55°C. DNA was separated on a 1% TBE-agarose gel and stained using SYBR Gold.

For phenol-chloroform partitioning of cleavage products, cleavage reactions were stopped in the presence or absent of proteinase K. Following 20-minutes incubation at 55°C, samples were mixed with an equal volume of phenol-chloroform-isoamyl alcohol and centrifuged for 5 minutes at 13000 rpm. The organic phase and interphase were back-extracted twice with 100 mM TRIS-HCl pH 8, 1 mM EDTA, 200 mM NaCl. The organic phase and aqueous phase were ethanol precipitated. DNA was resuspended in buffer containing 30 mM TRIS-HCl pH 8.5, 1 mM EDTA, 100 mM NaCl and 0.2 mg/ml proteinase K and incubated for 1 hour at 55°C. The DNA was again ethanol precipitated, resuspended in TE buffer, separated on a 1% TBE-agarose gel and stained using SYBR Gold.

For the analysis of DNA cleavage with linear substrates containing 0, 1, 3 or 6 copies of the Widom 601 sequence, fragments containing 1, 3 or 6 copies of Widom 601 were cloned into the multiple cloning site of pUC19 to yield pCCB1106, pCCB1107 and pCCB1108, respectively. The plasmids were PCR amplified with primers pl68 and vg001 (containing a 5′ 6-carboxyfluorescein dye) to yield linear substrates for the reaction. Cleavage reactions were performed in standard conditions with 500 nM ^MBP^SPO11 and 25 ng/µl linear fluorescent substrate). DNA was separated on a 1% TBE-agarose gel and visualized using a Typhoon scanner (Cytiva).

Nicked DNA substrates were prepared by treatment of pOC157 with Nb.BrsDI, followed by phenol extraction and ethanol precipitation.

### Detection of fluorescent SPO11-DNA covalent complexes

Substrates were assembled by annealing primers dd77 and cb100, or cb1593 and cb100, to produce 80 bp duplex DNA with a 6-FAM fluorophore located in 5′ or 3′, respectively. The oligos were mixed in equimolar concentrations (10 μM) in STE (100 mM NaCl, 10 mM Tris-HCl at pH 8, 1 mM EDTA), heated, and slowly cooled on a PCR thermocycler (3 min at 98°C, 1 hour at 75°C, 1 hour at 65°C, 30 min at 37°C, and 10 min at 25°C).

Cleavage reactions (20 µl) contained 1 µM ^MBP^SPO11, 0.5 µM fluorescent substrate in buffer containing 25 mM Tris pH 7.5, 5 % glycerol, 10 % DMSO, 40 mM NaCl, 1 mM DTT, 5 mM MgCl_2_ and 1.5 mM MnCl_2_. Reactions were incubated for 2 hours at 37°C, stopped with 1X Leammli buffer, and separated by SDS-PAGE. The fluorescent gel was scanned using a Typhoon scanner (Cytiva), and proteins stained with Coomassie blue.

### Sequencing gel analysis of SPO11 cleavage sites

Oligonucleotides cb95 and cb100 were first purified on 10% polyacrylamide-urea gels. For each oligo, 5 pmol were 5′-end-labelled with [γ-^32^P]-ATP and T4 polynucleotide kinase (NEB). The labeled oligo was mixed in equimolar concentrations with the unlabeled reverse complement and annealed by heating at 100°C in a water bath followed by slow cooling. Labeled substrates were then purified by native polyacrylamide gel electrophoresis.

Cleavage reactions (20 µl) contained 500 nM ^MBP^SPO11, 1 nM radioactive substrate and 2.5 nM (100 ng) plasmid DNA in buffer containing 25 mM Tris pH 7.5, 5 % glycerol, 0.1 mg/ml BSA, 50 mM NaCl, 1 mM DTT, 5 mM MgCl_2_ and 1.5mM MnCl_2_. Reactions were incubated for 2 hours at 37°C, stopped with 50 mM EDTA and 1% SDS. Markers were generated by partial digestion of the substrate using the indicated restriction enzymes or DNase I. The DNA was ethanol precipitated and separated on a 10% TBE-UREA sequencing gel. The gel was dried and developed by autoradiography.

### Gel shift assays

Binding reactions (20 μl) were carried out using the indicated concentration of protein with 100 ng plasmid DNA in buffer containing 25 mM Tris-HCl pH 7.5, 7.5% glycerol, 50 mM NaCl, 1 mM DTT, 5 mM EDTA, and 1 mg/ml BSA. Binding reactions were incubated for 30 minutes at 37°C and separated on 1% TAE-agarose gels at 60 V for 2 h. Gels were stained with SYBR Gold and scanned using a Typhoon scanner (Cytiva).

### End Matter

## Author Contributions and Notes

CCB conceived the project; CO and CCB secured funding, designed experiments, performed experiments, and analyzed data. CCB wrote the paper with input from CO. The authors declare no conflict of interest.

## Acknowledgments

We thank Scott Keeney for discussion and sharing unpublished information, and Bernard Hallet and Ronald Chalmers for comments on the manuscript. We thank CCB laboratory members Pascaline Liloku, Dima Daccache, Mahesh Survi and Théo Christopher Borremans for help with SEC-MALS experiments, insect cell maintenance, and protein purifications. This work was supported by the European Research Council under the European Union’s Horizon 2020 research and innovation program (ERC grant agreement 802525 to CCB). CO was funded in part by a fellowship from the Fonds National de la Recherche Scientifique (FNRS). CCB is a FNRS Research Associate.

---

**Extended Data Fig. 1:**
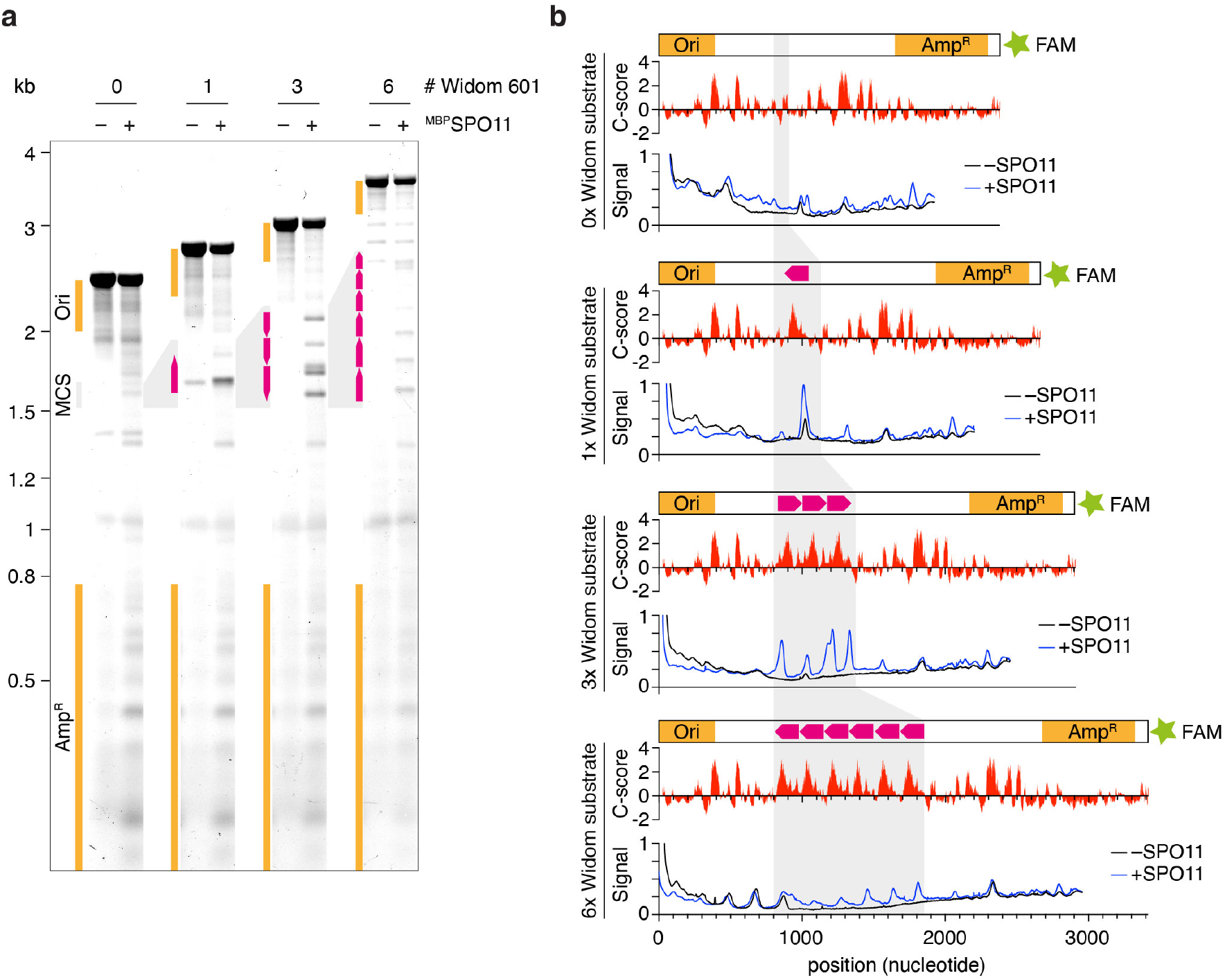
Correlation between DNA bendability and SPO11 activity. **(a)** Agarose gel analysis of SPO11 DNA cleavage with linear substrates containing 0, 1, 3, or 6 copies of the Widom 601 sequence. The pUC19-derived substrates contain a single fluorophore (FAM) at one end, allowing the mapping of cleavage products. The position of the origin of replication (Ori), ampicillin resistance gene (Amp^R^) and the multiple cloning site (MCS) are indicated. Copies of the Widom 601 sequence are represented with pink arrows. **(b)** Correlation between Widom 601 sequences, DNA bending, and SPO11 cleavage. The C-score is a bendability parameter predicted by DNAcycP^20^. The insertion of Widom 601 sequences creates hotspots for SPO11, although the predicted bendability of the DNA sequence is not sufficient to account for the cleavage activity observed along the substrate. The cleavage hotspots produced by the Widom 601 sequence can also be due to a sequence preference of SPO11, irrespective of bendability.

**Extended Data Fig. 2:**
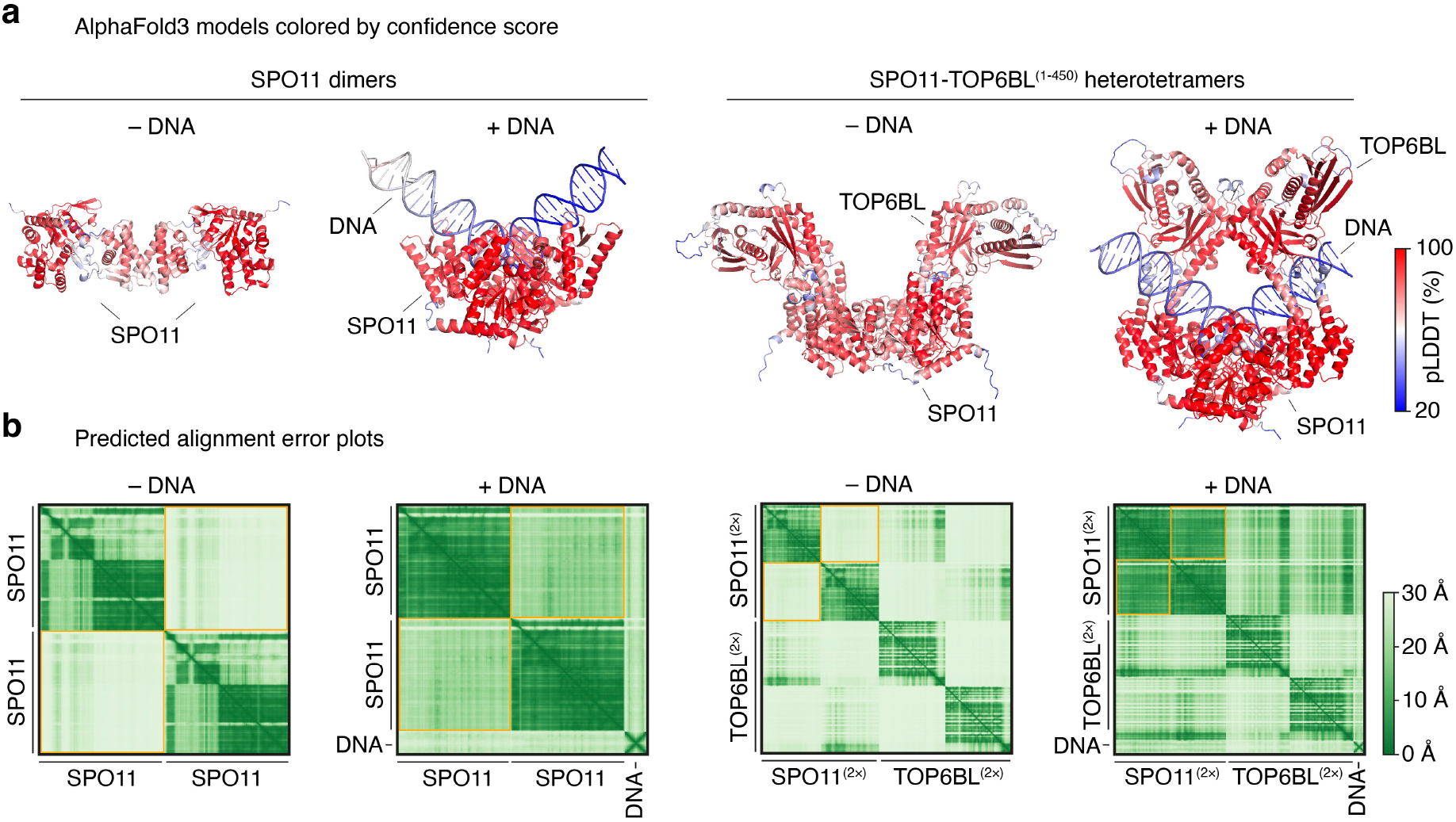
Quality assessment of AlphaFold3 models. **(a)** AlphaFold3 models colored by confidence score of SPO11 dimers and SPO11-TOP6BL complexes with or without 40 bp duplex DNA substrate. The C-terminus of TOP6BL was omitted from the model because it is predicted to be unstructured. **(b)** Predicted alignment error plots. The structure and relative position (orange squares) of SPO11 monomers are predicted with lower confidence in the absence of DNA than in the presence of DNA, consistent with the monomeric stoichiometry of SPO11 and SPO11-TOP6BL complexes. In the absence of DNA, AlphaFold proposes an aberrant dimeric model of SPO11 throught interactions between WH domains (left). In the presence of DNA and TOP6BL, the relative position of SPO11 is predicted with much higher accuracy (compare orange squares on the rightmost PAE plot with the others).

**Extended Data Fig. 3:**
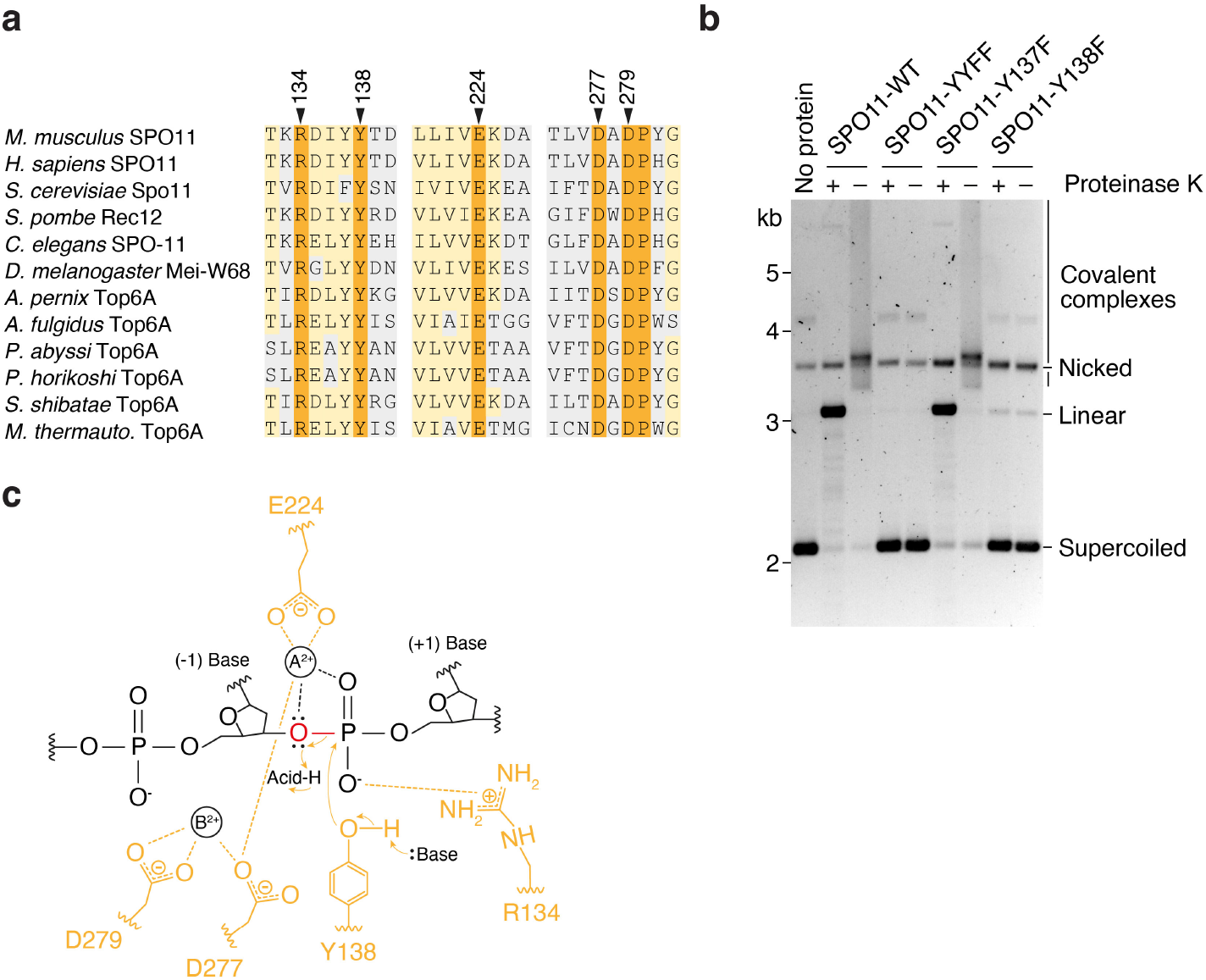
Two metal-ion mechanism and active site residues of SPO11. **(a)** Sequence alignments of eukaryotic SPO11 and archaeal Top6A proteins. Blocks around active site residues are shown. Invariant amino acids are in orange, conservative substitutions are in light orange. Active site residues are indicated with an arrowhead. **(b)** Cleavage assay with wild-type SPO11, double Y137F/Y138F (YYFF) and single Y137F and Y138F mutants. Conversion of the cleaved linear product into a smear in the absence of proteinase K indicates covalent attachment of SPO11 to the broken DNA ends. The low amount of linear and nicked products observed with the Y138F mutant is due to a non-specific nuclease contaminant in the protein preparation. **(c)** Two-metal-ion reaction scheme, based on the mechanism proposed for type IIA topoisomerases^26^.

**Extended Data Fig. 4:**
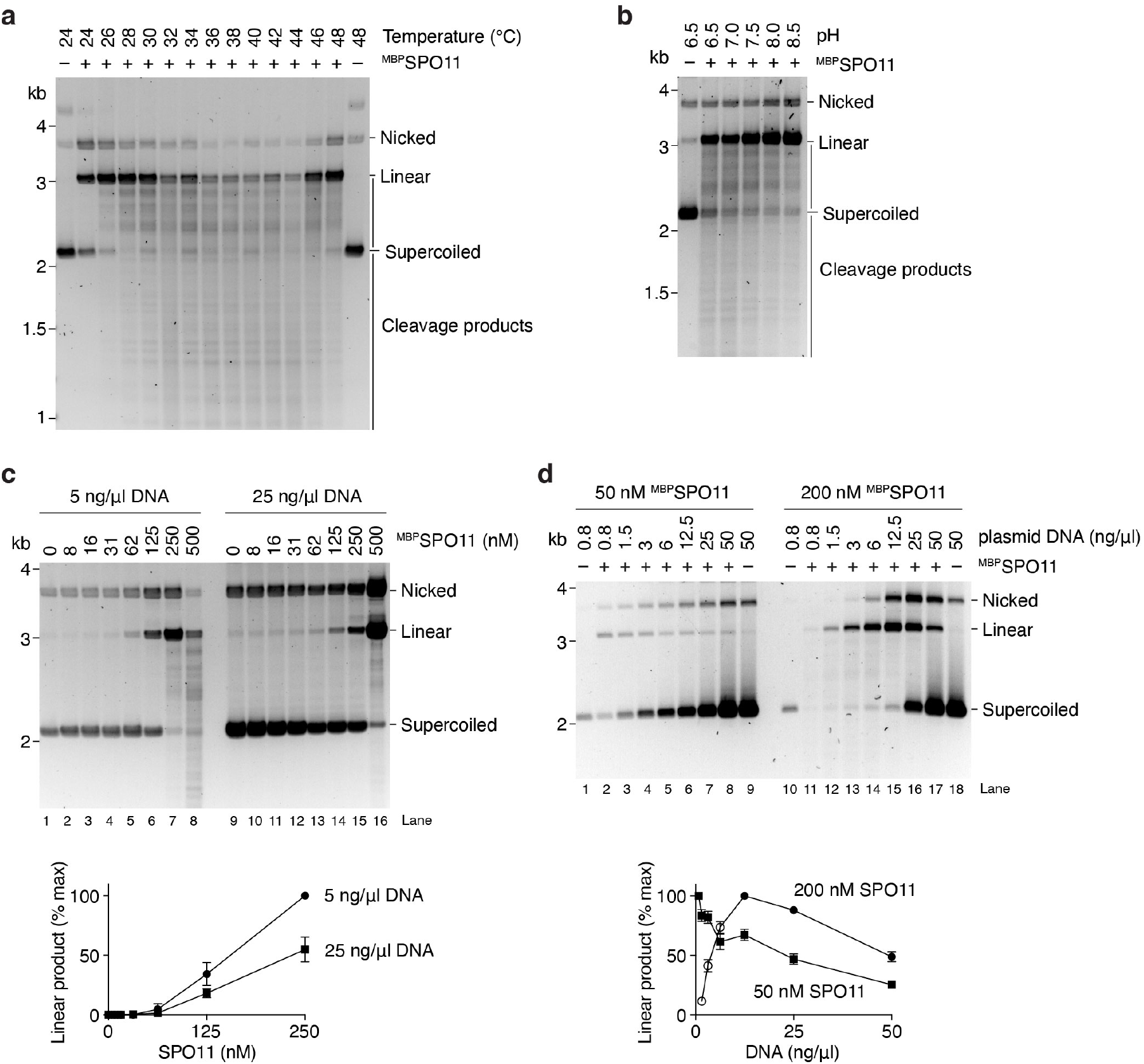
Optimization of the cleavage reaction. **(a, b)** Effect of the reaction temperature (a) and pH (b) on SPO11 cleavage. The standard conditions chosen for all the reactions are 37°C and pH 7.5. **(c)** Titration of SPO11 protein in reactions that contained either 5 ng/µl (2.5 nM) or 25 ng/µl (12.5 nM) plasmid DNA. Quantifications show the mean and range from two independent experiments. Cleavage increased with protein concentration, but the total level of cleavage was higher in reactions that had lower DNA concentrations (compare linear product in lanes 5-7 with lanes 13-15). **(d)** Titration of DNA in reactions that contained either 50 or 200 nM SPO11. At 50 nM SPO11, cleavage was highest at the lowest DNA concentration tested (0.8 ng/µl, 0.4 nM). At 200 nM SPO11, a reduction of total DNA cleavage was observed at substrate concentrations above 12.5 ng/µl (6.2 nM). At lower concentrations (open circles), the substrate is limiting so the amount of linear product is not representative of total break levels. The reaction time in these experiments is 2 hours. Note that in **Fig. 4d**, the inhibitory effect is observed at lower DNA concentrations because the reaction time was shorter (15 minutes). Longer reaction times provide more opportunity for dimerization and cleavage.

**Extended Data Fig. 5:**
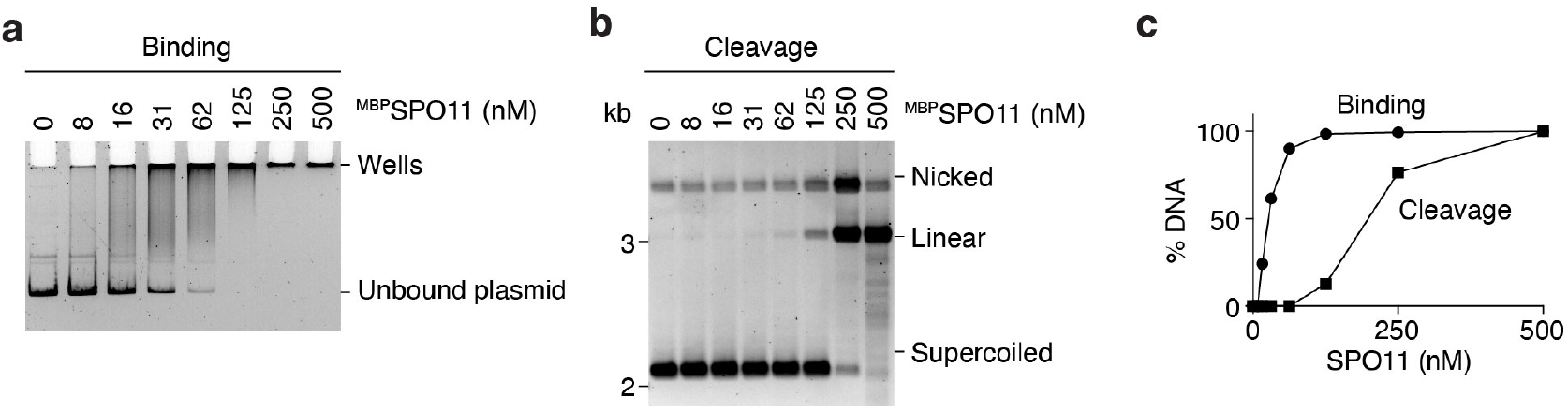
DNA cleavage requires higher SPO11 concentration than DNA binding. Comparison of the DNA-binding **(a)** and DNA cleavage **(b)** activities of SPO11. Reactions contained 5 ng/µl (2.5 nM) plasmid. Binding reactions were assembled for 30 minutes. Cleavage reactions were stopped after 2 hours. **(c)** Quantification shows that the plasmid is bound efficiently at concentrations that do not support cleavage.

**Extended Data Fig. 6:**
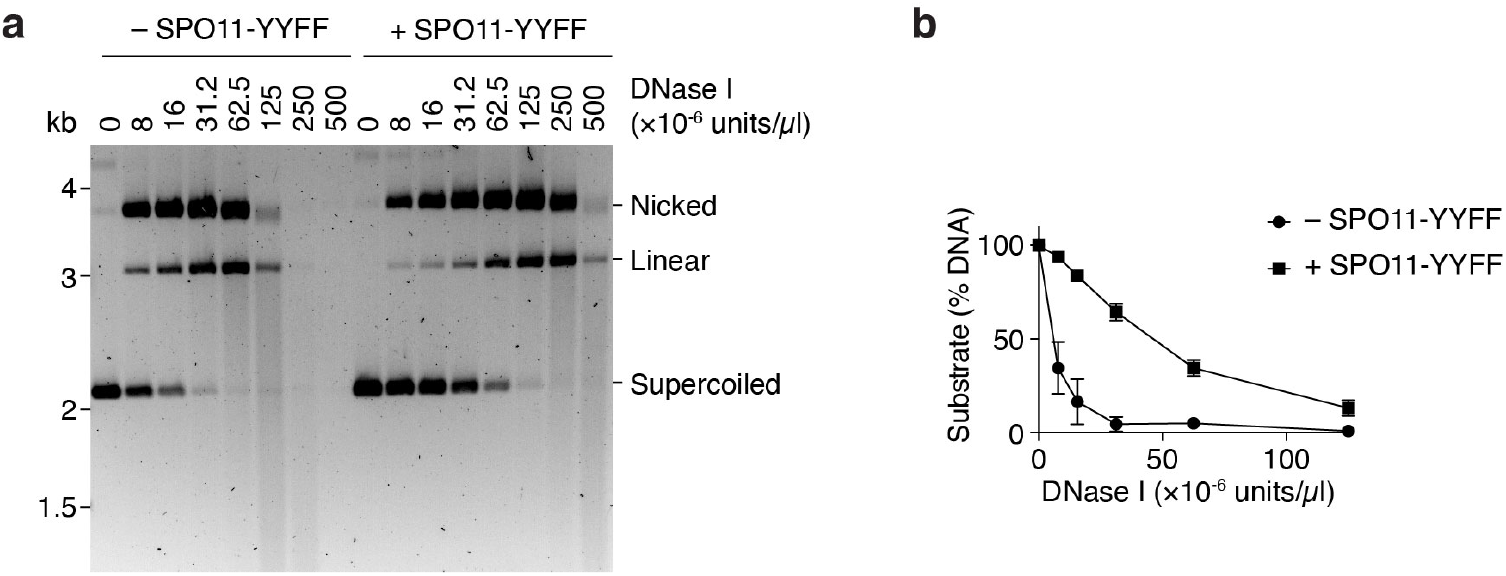
SPO11 provides effective protection against DNase I treatment. **(a)** Plasmid substrates (2.5 nM) were incubated with or without 600 nM catalytically inactive SPO11-YYFF mutant, followed by two-minute treatment with the indicated concentration of DNase I. **(b)** Quantification of the supercoiled substrate remaining in panel a. Error bars are ranges from two independent experiments.

**Extended Data Fig. 7:**
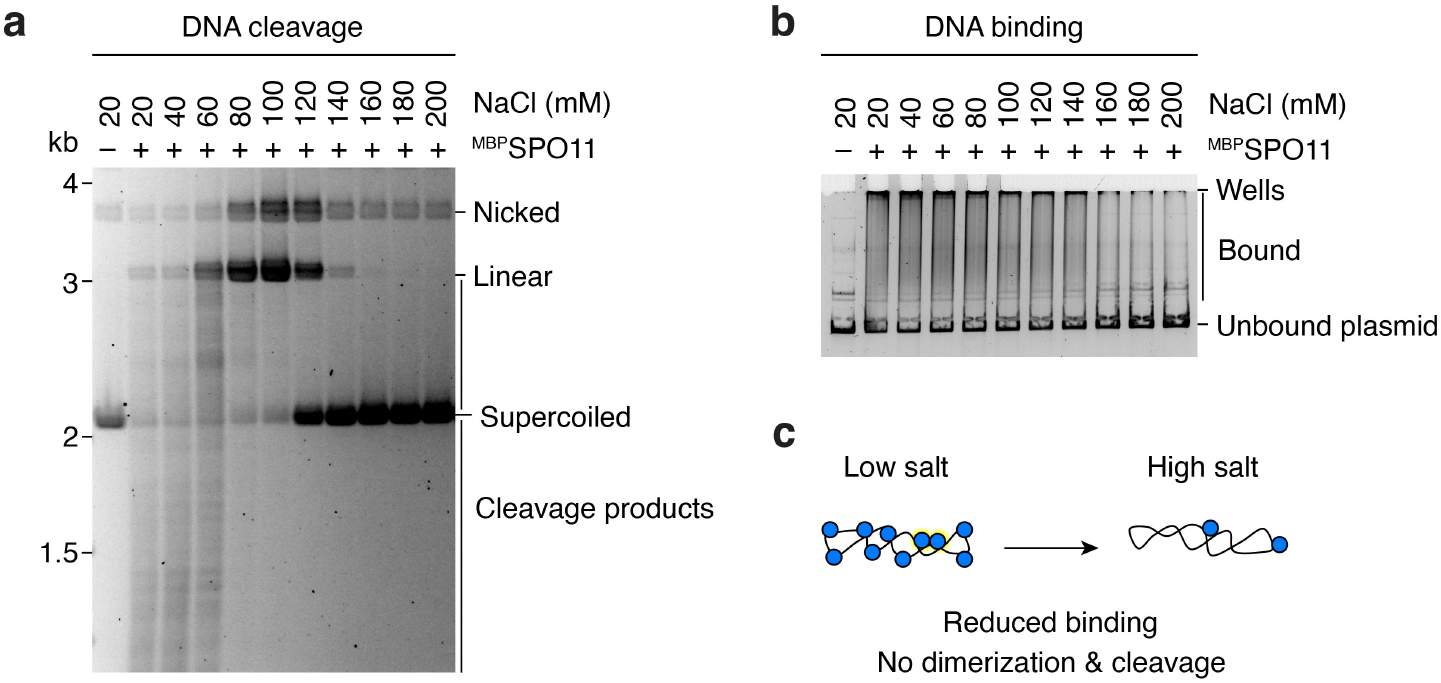
DNA binding and cleavage by SPO11 are sensitive to salt. Effect of NaCl on **(a)** plasmid cleavage and **(b)** DNA binding. SPO11 concentration is 500 nM in panel A and 100 nM in panel B. **(c)** The greater sensitivity of cleavage than DNA binding to salt could be explained by a mild reduction in DNA binding severely reducing the chances of dimerization on DNA.

**Extended Data Fig. 8:**
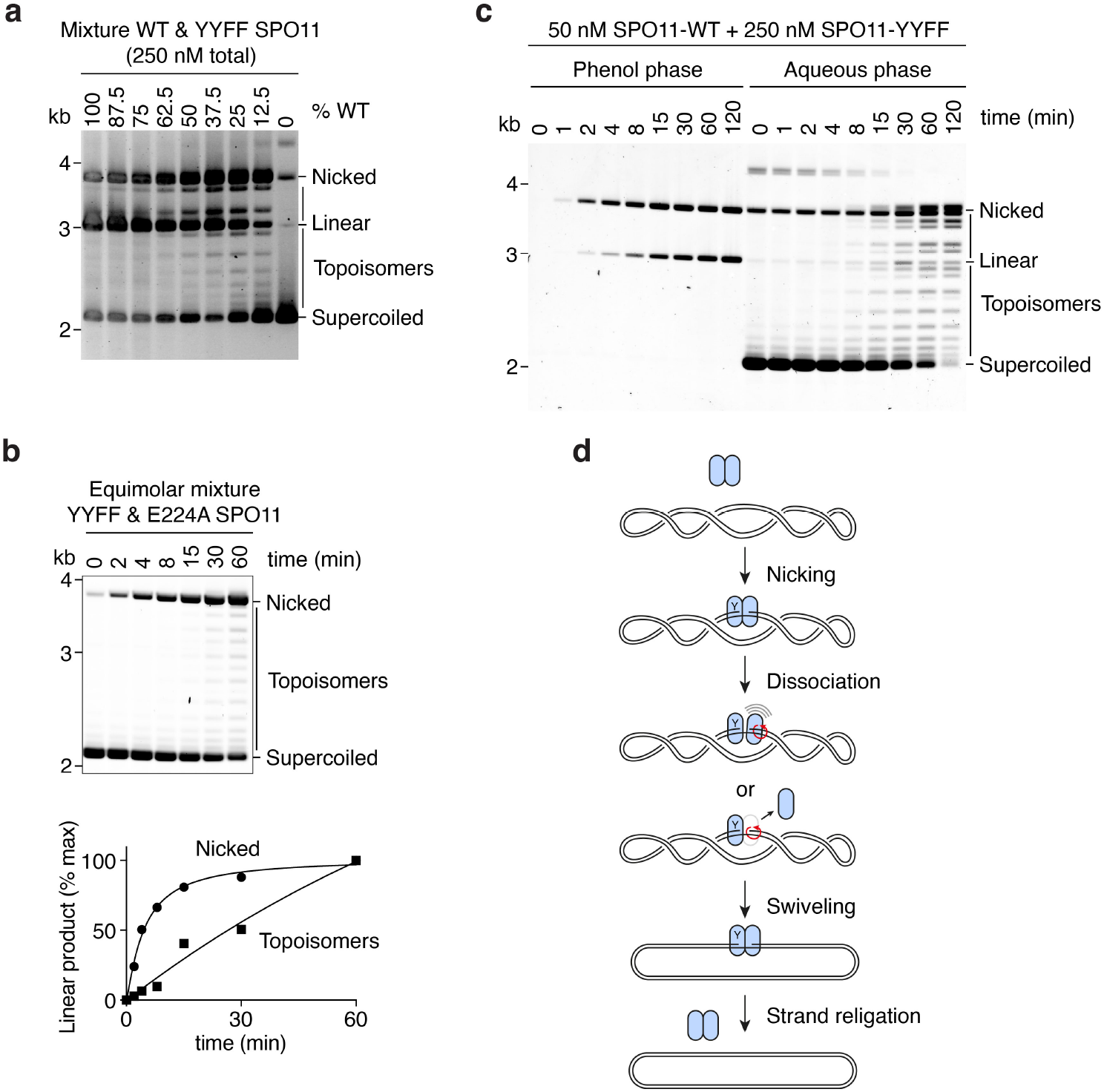
SPO11 can reseal single-strand DNA nicks. **(a)** Cleavage analysis at a constant SPO11 concentration in the presence of different ratios of wild-type and catalytically-inactive (YYFF) mutants. The ladder that migrates between the supercoiled and nicked products is absent in reactions that contain only wild-type or inactive mutants, indicating that it is not due to a contaminating activity in one of the protein preparations. **(b)** Kinetic analysis of cleavage products in reactions that contained mixtures of YYFF and E224A mutants. **(c)** Phenol-chloroform partitioning of DNA cleavage products in reactions that contained mixtures of wild-type and mutant SPO11. Topoisomers partition to the organic phase as the covalent link with SPO11 has been released. **(d)** Illustration of the plasmid relaxation activity observed with mixtures of wild type and inactive SPO11. Dissociation of the SPO11 dimer after single-strand nicking will lead to the swiveling of the DNA duplex around the intact phosphodiester bond (red arrows). Restauration of the SPO11 dimer then provides an opportunity for strand religation. Separation of the dimer interface could also be accompanied by the dissociation of the subunit not involved in catalysis from the DNA substrate, although this would be expected to lead to full plasmid relaxation, which is not always observed.

**Extended Data Fig. 9:**
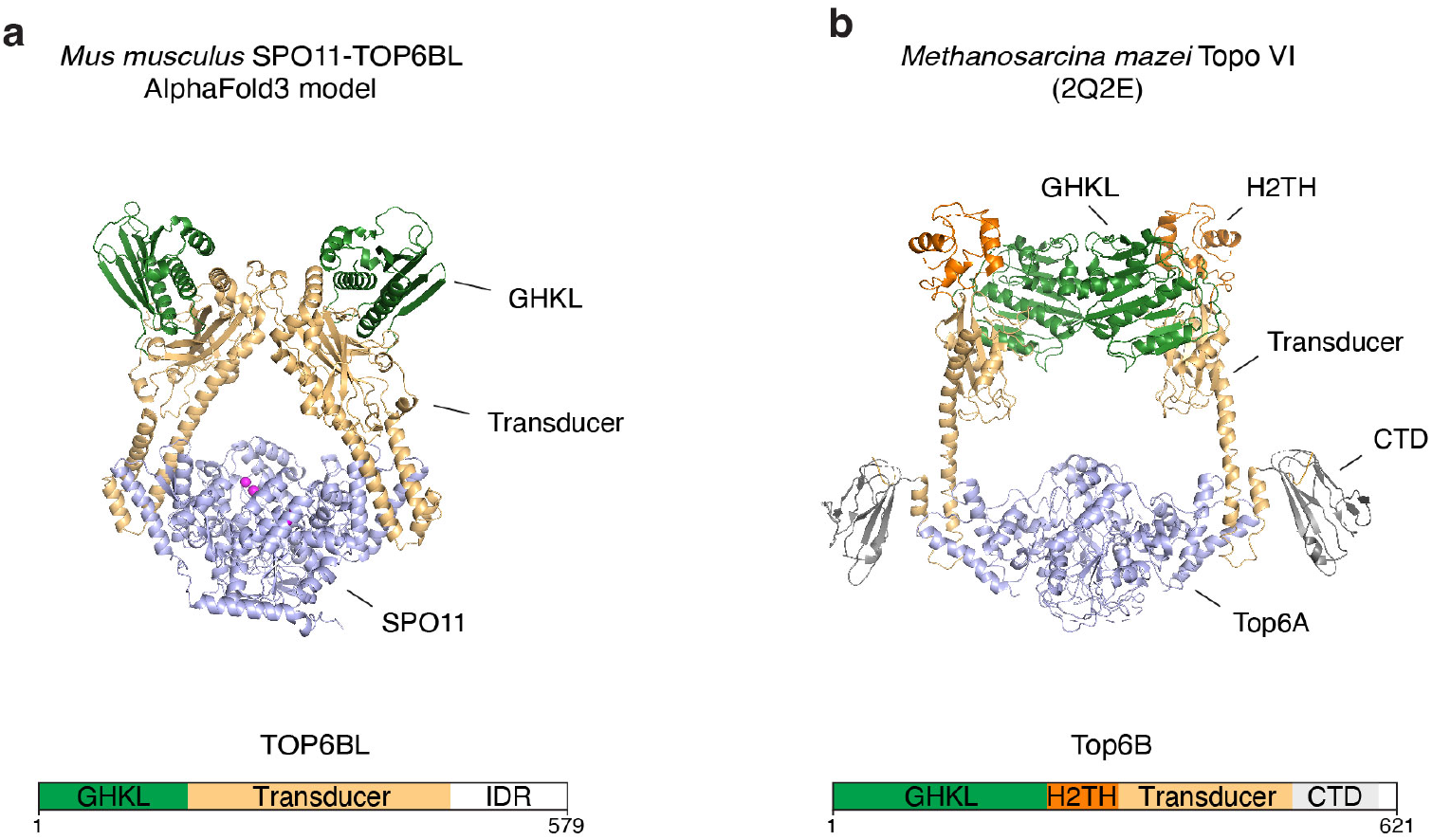
Comparison of the AlphaFold model of mouse SPO11-TOP6BL and the structure of *Methanosarcina mazei* Topo VI. **(a)** AlphaFold3 model of a 2:2 SPO11-TOP6BL heterotetramer. The structure was modeled with DNA, but the DNA was hidden to ease comparison. **(b)** Crystal structure of Topo VI^10^. SPO11 and Top6A are in blue. GHKL (green), helix two-turn helix (H2TH, orange), transducer (yellow), C-terminal domain (CTD, grey), intrinsically-disordered region (IDR, not shown).

**Supplementary Table S1:** Plasmids used in this study.

| Plasmid name | Description |
| --- | --- |
| pCCB628 | HisFlag-TOP6BL in pFastBac1 |
| pCCB630 | MBP-SPO11 in pFastBac1 |
| pCCB642 | MBP-SPO11(Y137F,Y138F) in pFastBac1 |
| pCCB1084 | MBP-SPO11(Y137F) in pFastBac1 |
| pCCB1085 | MBP-SPO11(Y138F) in pFastBac1 |
| pCCB1086 | MBP-SPO11(E224A) in pFastBac1 |
| pCCB959 | Standard 3-kb substrate for cleavage assay |
| pOC157 | Substrate with 24× Widom 601 sequences<br>(pBS 24x-601 MA1+MA2, Addgene plasmid #114361) |
| pUC19 | Control substrate with 0× Widom 601 |
| pCCB1106 | pUC19 with 1× Widom 601 |
| pCCB1107 | pUC19 with 3× Widom 601 |
| pCCB1108 | pUC19 with 6× Widom 601 |

**Supplementary Table S2:**
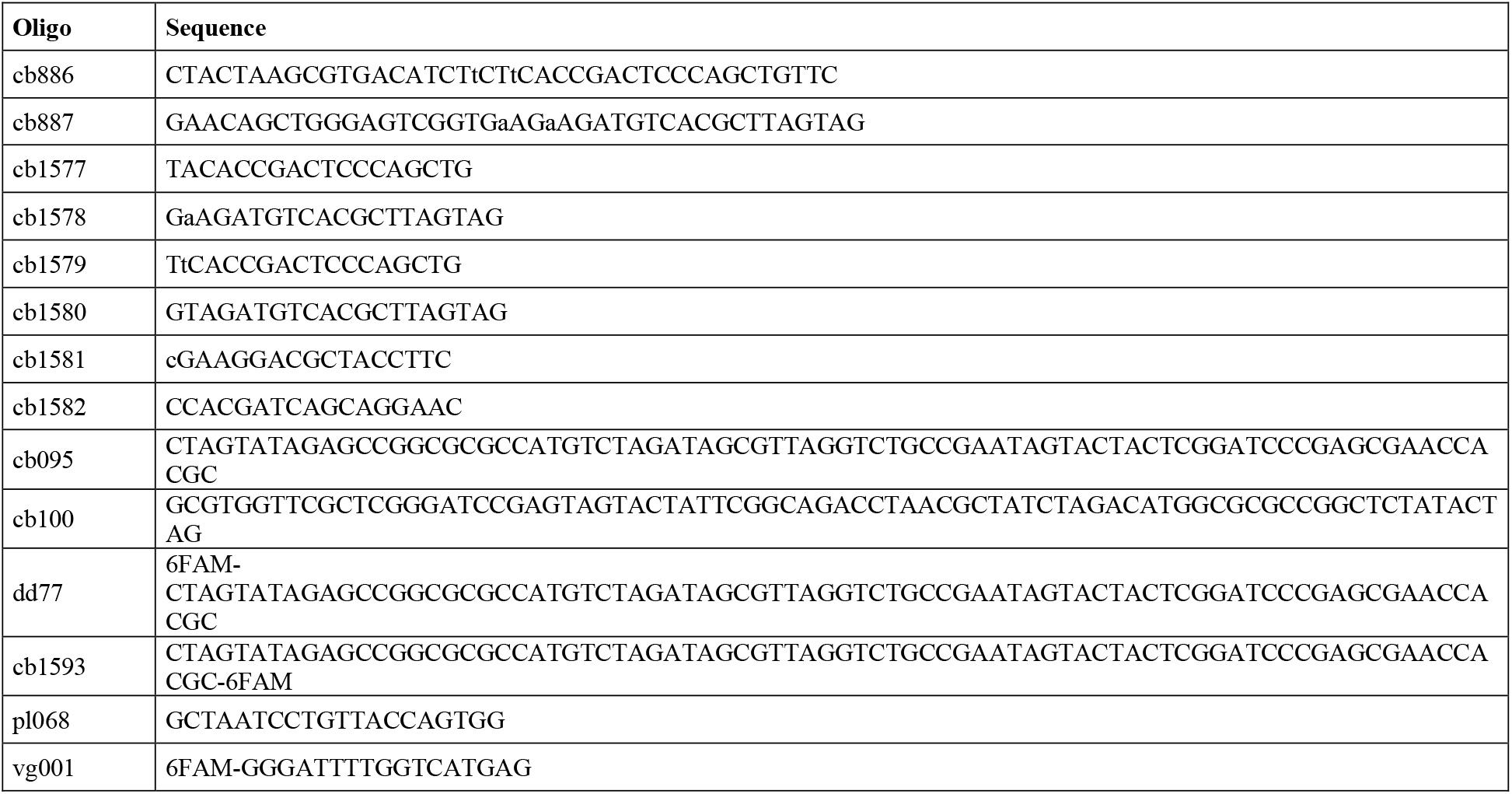
Oligonucleotides used in this study.

**Supplementary Table S3:**
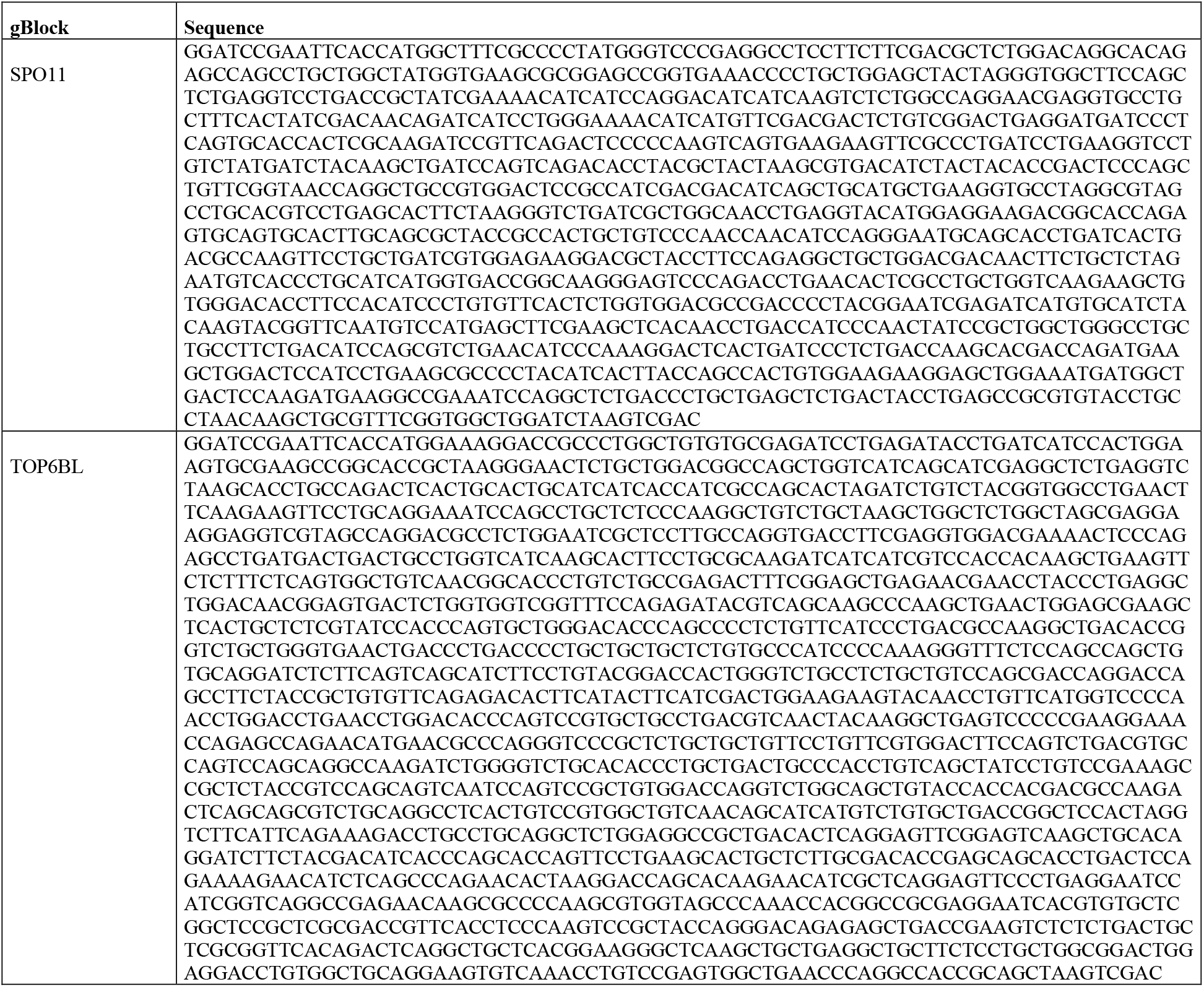
Synthetic genes used in this study.

## References

1 Keeney, S., Giroux, C. N. & Kleckner, N. Meiosis-specific DNA double-strand breaks are catalyzed by Spo11, a member of a widely conserved protein family. Cell 88, 375–384, doi:S0092-8674(00)81876-0 [pii] (1997).

2 Bergerat, A. et al. An atypical topoisomerase II from Archaea with implications for meiotic recombination. Nature 386, 414–417, doi:10.1038/386414a0 (1997).

3 de Massy, B. Initiation of meiotic recombination: how and where? Conservation and specificities among eukaryotes. Annu Rev Genet 47, 563–599, doi:10.1146/annurev-genet-110711-155423 (2013).

4 Robert, T., Vrielynck, N., Mezard, C., de Massy, B. & Grelon, M. A new light on the meiotic DSB catalytic complex. Semin Cell Dev Biol 54, 165–176, doi:10.1016/j.semcdb.2016.02.025S1084-9521(16)30065-9[pii] (2016).

5 Keeney, S. & Kleckner, N. Covalent protein-DNA complexes at the 5’ strand termini of meiosis-specific double-strand breaks in yeast. Proc Natl Acad Sci U S A 92, 11274–11278 (1995).

6 Liu, J., Wu, T. C. & Lichten, M. The location and structure of double-strand DNA breaks induced during yeast meiosis: evidence for a covalently linked DNA-protein intermediate. Embo j 14, 4599–4608 (1995).

7 Buhler, C., Lebbink, J. H., Bocs, C., Ladenstein, R. & Forterre, P. DNA topoisomerase VI generates ATP-dependent double-strand breaks with two-nucleotide overhangs. J Biol Chem 276, 37215–37222, doi:10.1074/jbc.M101823200M101823200 [pii] (2001).

8 Nichols, M. D., DeAngelis, K., Keck, J. L. & Berger, J. M. Structure and function of an archaeal topoisomerase VI subunit with homology to the meiotic recombination factor Spo11. Embo j 18, 6177–6188, doi:10.1093/emboj/18.21.6177 (1999).

9 Diaz, R. L., Alcid, A. D., Berger, J. M. & Keeney, S. Identification of residues in yeast Spo11p critical for meiotic DNA double-strand break formation. Mol Cell Biol 22, 1106–1115 (2002).

10 Corbett, K. D., Benedetti, P. & Berger, J. M. Holoenzyme assembly and ATP-mediated conformational dynamics of topoisomerase VI. Nat Struct Mol Biol 14, 611–619, doi:nsmb1264 [pii] ISSN 10.1038/nsmb1264 (2007).

11 Robert, T. et al. The Topo VIB-Like protein family is required for meiotic DNA double-strand break formation. Science 351, 943–949, doi:10.1126/science.aad5309 (2016).

12 Libby, B. J., Reinholdt, L. G. & Schimenti, J. C. Positional cloning and characterization of Mei1, a vertebrate-specific gene required for normal meiotic chromosome synapsis in mice. Proc Natl Acad Sci U S A 100, 15706–15711, doi:10.1073/pnas.2432067100 (2003).

13 Kumar, R., Bourbon, H. M. & de Massy, B. Functional conservation of Mei4 for meiotic DNA double-strand break formation from yeasts to mice. Genes Dev 24, 1266–1280, doi:10.1101/gad.571710 (2010).

14 Stanzione, M. et al. Meiotic DNA break formation requires the unsynapsed chromosome axis-binding protein IHO1 (CCDC36) in mice. Nat Cell Biol 18, 1208–1220, doi:10.1038/ncb3417 (2016).

15 Kumar, R. et al. Mouse REC114 is essential for meiotic DNA double-strand break formation and forms a complex with MEI4. Life science alliance 1, e201800259, doi:10.26508/lsa.201800259 (2018).

16 Baudat, F. et al. PRDM9 is a major determinant of meiotic recombination hotspots in humans and mice. Science 327, 836–840, doi:science.1183439 [pii] 10.1126/science.1183439 (2010).

17 Myers, S. et al. Drive against hotspot motifs in primates implicates the PRDM9 gene in meiotic recombination. Science 327, 876–879, doi:10.1126/science.1182363 (2010).

18 Parvanov, E. D., Petkov, P. M. & Paigen, K. Prdm9 controls activation of mammalian recombination hotspots. Science 327, 835, doi:10.1126/science.1181495 (2010).

19 Lange, J. et al. The Landscape of Mouse Meiotic Double-Strand Break Formation, Processing, and Repair. Cell 167, 695-708.e616, doi:10.1016/j.cell.2016.09.035 (2016).

20 Li, K., Carroll, M., Vafabakhsh, R., Wang, X. A. & Wang, J. P. DNAcycP: a deep learning tool for DNA cyclizability prediction. Nucleic Acids Res 50, 3142–3154, doi:10.1093/nar/gkac162 (2022).

21 Lowary, P. T. & Widom, J. New DNA sequence rules for high affinity binding to histone octamer and sequence-directed nucleosome positioning. J Mol Biol 276, 19–42, doi:10.1006/jmbi.1997.1494 (1998).

22 Berger, J. M. Type II DNA topoisomerases. Curr Opin Struct Biol 8, 26–32, doi:10.1016/s0959-440x(98)80006-7 (1998).

23 Champoux, J. J. DNA topoisomerases: structure, function, and mechanism. Annu Rev Biochem 70, 369–413, doi:10.1146/annurev.biochem.70.1.369 (2001).

24 Keeney, S. Spo11 and the Formation of DNA Double-Strand Breaks in Meiosis. Genome Dyn Stab 2, 81–123, doi:10.1007/7050_2007_026 (2008).

25 Schmidt, B. H., Burgin, A. B., Deweese, J. E., Osheroff, N. & Berger, J. M. A novel and unified two-metal mechanism for DNA cleavage by type II and IA topoisomerases. Nature 465, 641–644, doi:10.1038/nature08974 (2010).

26 Deweese, J. E., Burgin, A. B. & Osheroff, N. Human topoisomerase IIalpha uses a two-metal-ion mechanism for DNA cleavage. Nucleic Acids Res 36, 4883–4893, doi:10.1093/nar/gkn466 (2008).

27 Claeys Bouuaert, C. et al. Structural and functional characterization of the Spo11 core complex. Nat Struct Mol Biol, doi:10.1038/s41594-020-00534-w (2021).

28 Yeh, H. Y., Lin, S. W., Wu, Y. C., Chan, N. L. & Chi, P. Functional characterization of the meiosis-specific DNA double-strand break inducing factor SPO-11 from C. elegans. Sci Rep 7, 2370, doi:10.1038/s41598-017-02641-z (2017).

29 Nore, A. et al. TOPOVIBL-REC114 interaction regulates meiotic DNA double-strand breaks. Nat Commun 13, 7048, doi:10.1038/s41467-022-34799-0 (2022).

30 Diagouraga, B. et al. The TOPOVIBL meiotic DSB formation protein: new insights from its biochemical and structural characterization. Nucleic Acids Res, doi:10.1093/nar/gkae587 (2024).

31 Wendorff, T. J. & Berger, J. M. Topoisomerase VI senses and exploits both DNA crossings and bends to facilitate strand passage. Elife 7, doi:10.7554/eLife.31724 (2018).

32 Claeys Bouuaert, C. et al. DNA-driven condensation assembles the meiotic DNA break machinery. Nature 592, 144–149, doi:10.1038/s41586-021-03374-w (2021).

33 Yu, Y. et al. Cryo-EM structure of the Spo11 core complex bound to DNA. bioRxiv, 2023.2010.2031.564985, doi:10.1101/2023.10.31.564985 (2023).

34 Yadav, V. K. & Claeys Bouuaert, C. Mechanism and Control of Meiotic DNA Double-Strand Break Formation in S. cerevisiae. Frontiers in cell and developmental biology 9, 642737, doi:10.3389/fcell.2021.642737 (2021).

35 Lam, I. & Keeney, S. Mechanism and regulation of meiotic recombination initiation. Cold Spring Harb Perspect Biol 7, a016634, doi:10.1101/cshperspect.a016634 [pii] cshperspect.a016634 [pii] (2015).

36 Daccache, D. et al. Evolutionary conservation of the structure and function of meiotic Rec114-Mei4 and Mer2 complexes. Genes Dev 37, 535–553, doi:10.1101/gad.350462.123 (2023).

37 Liu, K. et al. Structure and DNA bridging activity of the essential Rec114– Mei4 trimer interface. bioRxiv, 2023.2001.2018.524603, doi:10.1101/2023.01.18.524603 (2023).

38 Johnson, D. et al. Concerted cutting by Spo11 illuminates meiotic DNA break mechanics. Nature 594, 572–576, doi:10.1038/s41586-021-03389-3 (2021).

39 Prieler, S. et al. Spo11 generates gaps through concerted cuts at sites of topological stress. Nature 594, 577–582, doi:10.1038/s41586-021-03632-x (2021).

40 Lukaszewicz, A., Lange, J., Keeney, S. & Jasin, M. De novo deletions and duplications at recombination hotspots in mouse germlines. Cell 184, 5970-5984.e5918, doi:10.1016/j.cell.2021.10.025 (2021).

41 Tisi, R., Vertemara, J., Zampella, G. & Longhese, M. P. Functional and structural insights into the MRX/MRN complex, a key player in recognition and repair of DNA double-strand breaks. Comput Struct Biotechnol J 18, 1137–1152, doi:10.1016/j.csbj.2020.05.013 (2020).

